# Early divergence and gene exchange highways in the evolutionary history of *Mesoaciditogales*

**DOI:** 10.1101/2023.06.22.546130

**Authors:** Anne A. Farrell, Camilla L. Nesbø, Olga Zhaxybayeva

## Abstract

The placement of a non-hyperthermophilic order *Mesoaciditogales* at the base of *Thermotogota* tree challenges the prevailing hypothesis that the last common ancestor of *Thermotogota* was a hyperthermophile. Yet, given the long branch leading to the only two *Mesoaciditogales* described to-date, the phylogenetic position of the order may be due to the long branch attraction artifact. By testing various models and applying data recoding in phylogenetic reconstructions, we observed that *Mesoaciditogales*’ basal placement is strongly supported by the conserved marker genes assumed to be vertically inherited. However, based on the taxonomic content of 1,181 gene families and a phylogenetic analysis of 721 gene family trees, we also found that a substantial number of *Mesoaciditogales* genes are more closely related to species from the order *Petrotogales*. These genes contribute to coenzyme transport and metabolism, fatty acid biosynthesis, genes known to respond to heat and cold stressors, and include many genes of unknown functions. The *Petrotogales* comprise moderately thermophilic and mesophilic species with similar temperature tolerances to that of *Mesoaciditogales*. Our findings hint at extensive horizontal gene transfer between, or parallel independent gene gains by, the two ecologically similar lineages, and suggest that the exchanged genes may be important for adaptation to comparable temperature niches.

**Significance:** The high-temperature phenotype is often referenced when conjecturing about characteristics of the last common ancestor of all present-day organisms. Such inferences rely on accuracy of phylogenetic trees, especially with respect to lineages that branch closest to the last common ancestor. Here, we examined evolutionary history of *Mesoaciditogales*, an early-branching lineage within *Thermotogota* phylum, which is one of the early-diverging groups of bacteria. *Thermotogota* is composed of thermophiles, hyperthermophiles and mesophiles, who collectively can grow between 20 to 90 degrees Celsius, making it challenging to infer the growth temperature of their common ancestor. Our analysis revealed a complex evolutionary history of *Mesoaciditogales’* genome content impacted by horizontal gene transfer, highlighting the challenges of ancestral phenotype inferences using present-day genomes.

## Introduction

The *Thermotogota* is a bacterial phylum encompassing thermophiles, hyperthermophiles and mesophiles, with their optimum growth temperatures (OGTs) ranging from 37 to 80°C. Since the first isolated members of the phylum were hyperthermophiles (Patel et al. 1985; Huber et al. 1986; Antoine et al. 1997), the extant *Thermotogota* species have long been assumed to have descended from a hyperthermophilic last common ancestor (LCA) (Zhaxybayeva et al. 2009; Butzin et al. 2013; Green et al. 2013). Later discoveries of thermophilic and mesophilic members of the phylum were inferred to be secondary adaptations of some *Thermotogota* members to lower OGTs (Pollo et al. 2015).

In 2013, a new *Thermotogota* member was identified - the moderate thermophile *Mesoaciditoga lauensis* and the 16S rRNA phylogeny placed this novel organism as an early-diverging taxon of the phylum (Reysenbach et al. 2013). Shortly after, a sister taxon of *M. lauensis*, *Athalassotoga saccharophila*, was described (Itoh et al. 2016). Currently, these taxa are the only two characterized species of the *Mesoaciditogales* order. Most 16S rRNA phylogenetic trees place *Mesoaciditogales* basal to all other *Thermotogota* (Itoh et al. 2016; L’Haridon et al. 2019). The OGTs of *M. lauensis* and *A. saccharophila* are 57-60°C and 55°C (Itoh et al. 2016), respectively, which are far below the threshold of 80°C for hyperthermophily. As a result, *Mesoaciditogales*’ basal placement challenges inferences about the hyperthermophilic phenotype of the phylum’s LCA and existing hypotheses about how *Thermotogota* taxa have adapted to different growth temperatures. It also affects inferences about phenotype of the LCA of the *Bacteria* domain (Stetter 1996; Galtier et al. 1999; Brochier & Philippe 2002; Barion et al. 2007; Boussau et al. 2008; Catchpole & Forterre 2019).

Despite the basal placement in 16S rRNA phylogenies, *Mesoaciditogales* share some common features with a subset of later-diversified *Thermotogota*. First, *M. lauensis* and *A. saccharophila* can grow at temperatures as low as 45°C and 30°C, respectively (Reysenbach et al. 2013; Itoh et al. 2016), making their temperature tolerances similar to multiple species in the *Petrotogales* order, whose OGTs range from 37 to 70°C (Davey et al. 1993; L’Haridon et al. 2021; Wery et al. 2001). Second, outside of *Mesoaciditogales*, *Mesoaciditoga lauensis*’ 16S rRNA gene has the closest nucleotide identity to that of the *Kosmotoga* spp. (Reysenbach et al. 2013). Finally, in two recent 16S rRNA gene phylogenies, *Mesoaciditogales* branch within *Petrotogales* (Steinsbu et al. 2016; Mori et al. 2020).

Observation of conflicting relationships between *Mesoaciditogales* and the rest of *Thermotogota* in different analyses raises the possibility that the basal position of *Mesoaciditogales* is an artefact. There are two reasons to question either of the two inferred *Mesoaciditogales’* positions within Thermotogota. First, *Mesoaciditogales*’ 16S rRNA gene sequences are very divergent from the rest of *Thermotogota*, and therefore the basal position of *Mesoaciditogales* could be due to long branch attraction (Felsenstein 1978; Bergsten 2005).

Second, OGT affects GC content in rRNA stems (Galtier & Lobry 1997; Green et al. 2013) and amino acid composition of proteins (Zeldovich et al. 2007; Sauer & Wang 2019). As a result, standard phylogenetic models may not adequately account for heterogeneity in sequence composition of *Thermotogota*, which are adapted to a wide range of OGTs.

In this study, we investigate phylogenetic placement of *Mesoaciditogales* in relationship to other *Thermotogota* by analyzing (i) 16S rRNA gene sequences from *Thermotogota* genomes and direct environmental amplifications, and (ii) protein-coding genes from the genomes of *M. lauensis* and *A. saccharophila*, representative *Thermotogota* isolates and metagenome-assembled genomes (MAGs) of other *Thermotogota*. Our dataset is designed to break long branches leading to *M. lauensis* and *A. saccharophila*, and we use substitution models and data recoding tailored to deal with heterogeneous datasets. Our analyses support the basal position of *Mesoaciditogales* within *Thermotogota*, but also reveal that some parts of *Mesoaciditogales* genomes share evolutionary history with genomes of bacteria from *Petrotogales* order.

## Results

### 16S rRNA gene tree supports the basal placement of Mesoaciditogales within Thermotogota, but produces an unexpected position of Kosmotogaceae

To shorten the lengths of the *Mesoaciditogales* branches in comparison to the rest of the phylum members, we augmented the dataset of the 16S rRNA genes from 55 described *Thermotogota* species with the environmental 16S rRNA gene sequences most closely related to *Mesoaciditogales*. There is a substantial variation in GC content of *Thermotogota* 16S rRNA genes due to correlation of the GC-content of stems in the secondary structure with optimum growth temperature (Green et al. 2013). Indeed, among 277 sequences in our dataset, 23 out of 63 16S rRNA genes from the described species and 16 of the 203 environmental sequences failed the χ^2^ test of compositional homogeneity. The dataset as a whole also failed the Bowker’s test (Dutheil & Boussau 2008) under a homogenous model (Global Bowker’s test, p = 0.0019). These results suggest that the dataset is compositionally heterogeneous, raising the possibility of an incorrect phylogenetic inference under a homogeneous model. To address this issue, we reconstructed trees under both homogeneous and non-homogeneous models.

Phylogenies under both types of models have identical relations among the *Thermotogota* genera, and *Mesoaciditogales* clade is situated at the base of the *Thermotogota* phylum (**Supplementary Figure 1**). The addition of environmental sequences shortens the lengths of the *Mesoaciditogales* branches in comparison to the rest of the phylum members, reducing the possibility that the basal position of *Mesoaciditogales* is due to the long branch attraction artefact (Felsenstein 1978; Bergsten 2005). However, both trees exhibit a non-conventional relationship among *Thermotogota* families: the *Kosmotogaceae* family groups closest to *Mesoaciditogales* order instead of being a sister clade to the *Petrotogaceae* family. Notably, this relationship is different from the *Mesoaciditogales* relationships with *Petrotogales* observed in some 16S rRNA phylogenies (Steinsbu et al. 2016; Mori et al. 2020).

To further evaluate if there are specific taxa responsible for placement of *Kosmotogaceae* near *Mesoaciditogales*, we performed likelihood mapping analysis (Schmidt & Haeseler 2007) on an alignment that contained only described *Thermotogota* species and an outgroup.

Surprisingly, only 37.6% of quartets strongly support a relationship consistent with a basal position of *Mesoaciditogales*. The plurality of quartets (45.6%) strongly support grouping of *Mesoaciditogales* with *Petrotogales* (**Supplementary Figure 2A**). In an additional likelihood mapping analysis, aimed at evaluating if relationships between *Mesoaciditogales* and *Petrotogales* orders are due to specific relationship of *Mesoaciditogales* with either *Kosmotogaceae* or *Petrotogaceae* families, the plurality of quartets (45.7%) strongly support *Mesoaciditogales* grouping with *Petrotogaceae,* while only 14.8% of the quartets cluster *Mesoaciditogales* with *Kosmotogaceae* (**Supplementary Figure 2B**). These surprising relationships between *Mesoaciditogales* and *Petrotogales* are consistent with the observation that the 16S rRNA genes of *M. lauensis* and *A. saccharophila* are generally most similar to those of the described *Kosmotogaceae* (median 76.8% nt. identity) and *Petrotogaceae* (76.1%) species than to those of *Thermotogales* (74.8%), although there is an overlap of pairwise identities to specific members of these taxonomic groups (**Supplementary Tables 1** and **2**).

The *Mesoaciditogales* and *Petrotogales* relationships could be an artifact of nucleotide composition biases. Specifically, due to similarities in OGT between described species of *Mesoaciditogales* and *Petrotogales,* the GC content of their 16S rRNA genes (either for the full-length sequence or for stem regions only) is more similar to each other than to that of *Thermotogales* (**Supplementary Tables 3** and **4**). If the relationship between *Mesoaciditogales* and *Petrotogales* 16S rRNA genes is not an artifact, we should observe it in phylogenies of other genes. Therefore, we expanded our analyses to genes encoding ribosomal proteins and other broadly conserved proteins, which are widely used as universal markers to infer organismal relationships (Wolf et al. 2001; Gevers et al. 2005; Yutin et al. 2012).

### Trees reconstructed from ribosomal proteins and single copy core proteins also support the basal placement of *Mesoaciditogales*

The tree based on 50 concatenated ribosomal proteins from described *Thermotogota* species supports the deeply branching placement of *Mesoaciditogales* (**Supplementary Figure 3**). However, unlike 16S rRNA gene phylogenies, the topology shows the conventional placement of *Kosmotogaceae* as a sister group to *Petrotogaceae* (Nesbø et al. 2021). Since this tree can also suffer from the effects of long branch attraction artifacts, we added sequences from MAGs that broke multiple long branches, including the branch leading to *Mesoaciditogales*. No change to the topology was observed (**Supplementary Figure 4**). Individually, the majority of ribosomal proteins (33 out of 50) strongly support the basal position of *Mesoaciditogales*, and only 6 of 50 ribosomal proteins place *Mesoaciditogales* taxa with *Petrotogales* (**Supplementary Figure 5**).

Expanding the ribosomal protein families to 232 protein families encoded by single-copy core (SCC) genes, defined as genes present in at least 75% of described *Thermotogota* species, did not change the phylogeny (**Figure 1**). We further tested the strength of this phylogenetic signal by comparing the likelihoods of several alternative relationships between *Mesoaciditogales* and other *Thermotogota* families. All alternative topologies were rejected (an approximately unbiased [AU] test, p-value < 0.001; **Figure 2**).

**Figure 1.**
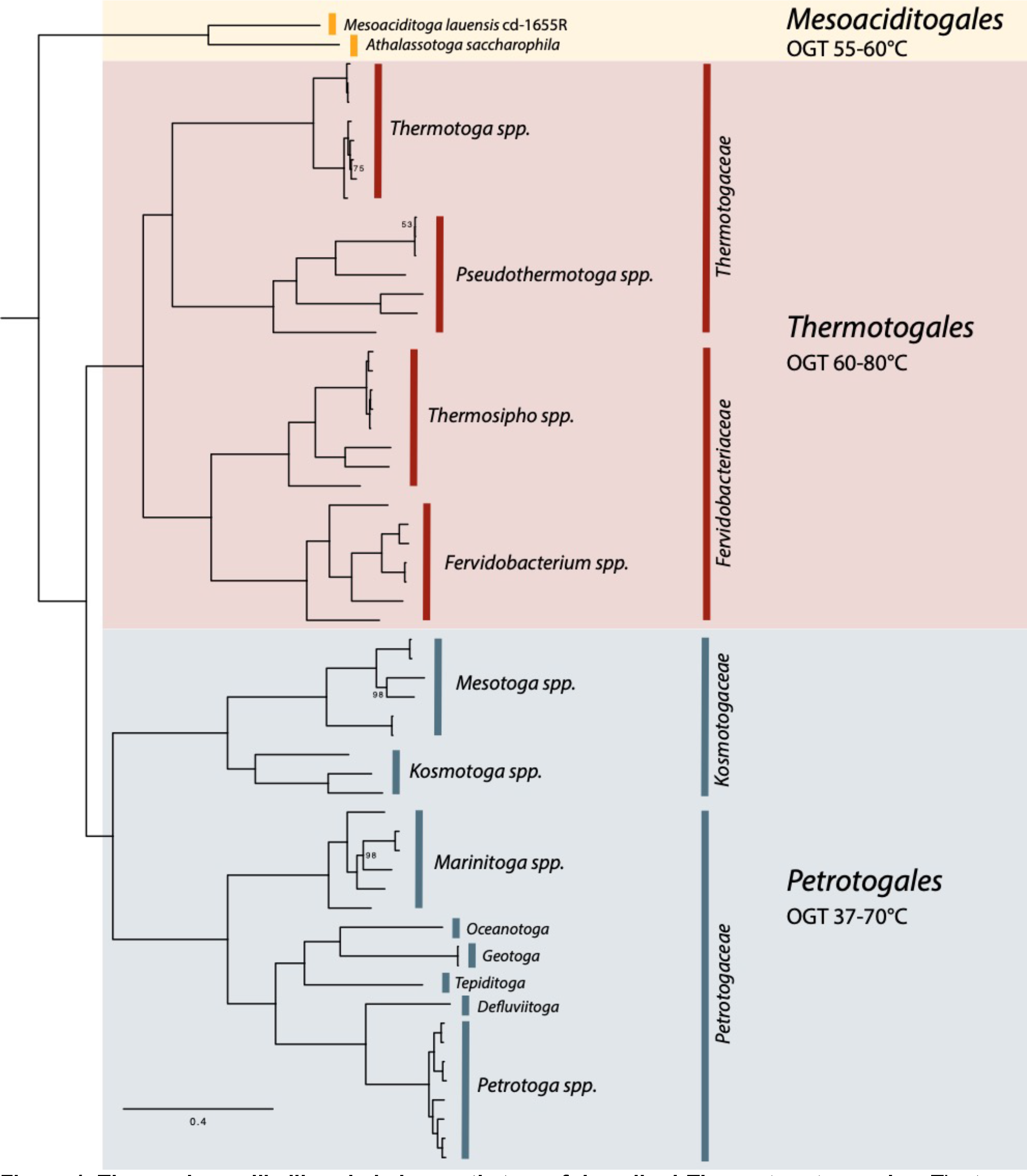
**The maximum likelihood phylogenetic tree of described *Thermotogota* species.** The tree was reconstructed from amino acid sequences of 232 single-copy core genes. Bootstrap support values below 100% are shown as values at the branches; all unlabeled branches have 100% bootstrap support. Scale bar, substitutions per site. The tree with taxa labels of individual branches is available in the **FigShare repository** (see **Methods**).

**Figure 2.**
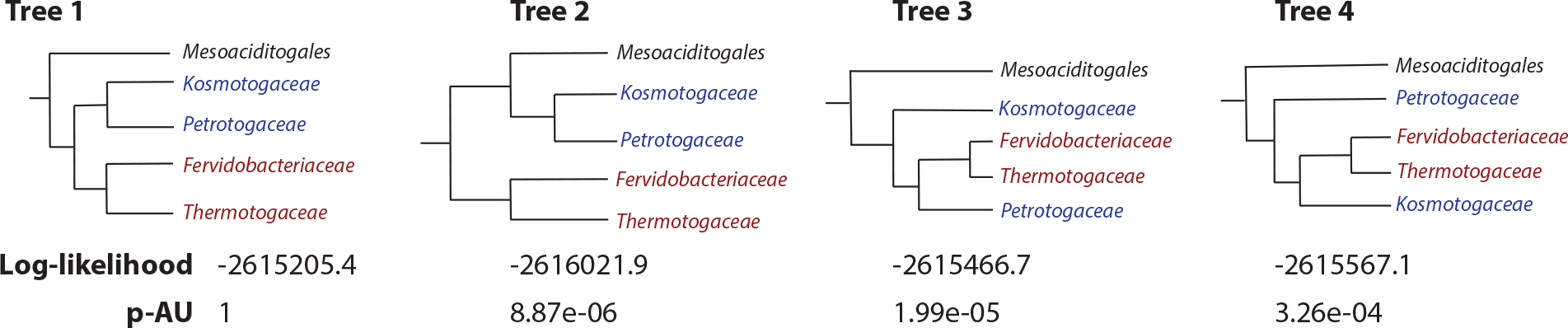
**Support of the four different relationships among *Thermotogota* families by the concatenated SCC dataset**. Tree 1 represents the relationships shown on Figure 1 and is the null hypothesis. The tested alternative relationships are shown as Tree 2, Tree 3 and Tree 4. “Log-Likelihood” is logarithm of likelihood of the SCC alignment given the tree. “p-AU” is the p-value of the AU test.

### Accounting for the amino acid composition bias of *Thermotogota* proteins does not affect basal position of *Mesoaciditogales*

Optimal growth temperature results not only in GC bias in 16S rRNA genes, but also in overrepresentation of certain amino acids in proteins, such as IVYWREL (Zeldovich et al. 2007), and these biases have been observed in *Thermotogota* (**Figure 3** and Zhaxybayeva et al. 2009; Nesbø et al. 2012; Pollo et al. 2015).

**Figure 3.**
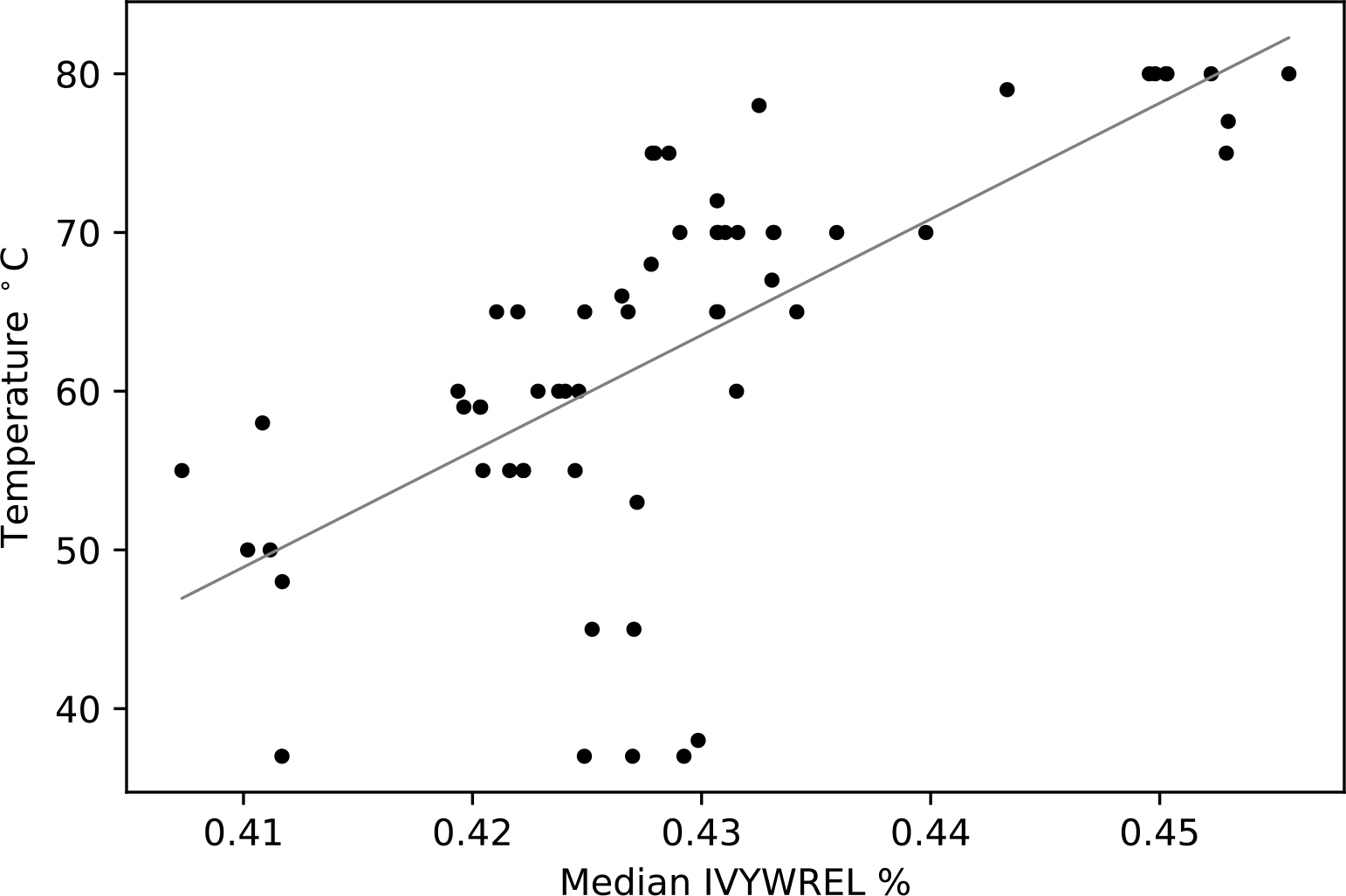
Correlation of OGT in described *Thermotogota* species with the median IVYWREL amino acid composition bias in proteins encoded in their genomes [R^2^ = 0.46, mean squared error = 78.06, *y = 731.0x -250.8*]. Each point represents a genome of a species. The median IVYWREL bias was calculated from IVYWREL biases of all cytosolic proteins encoded in a genome (see **Methods** for details).

The amino acid compositional biases are known to affect phylogenetic inferences that employ standard, homogeneous models (Foster & Hickey 1999; Li et al. 2014). Unfortunately, all sequences in the alignment of the concatenated SCC proteins failed IQ-TREE’s χ^2^ compositional heterogeneity test (p-value < 0.05; df=19), which is likely linked to the above-described amino acid bias.

To correct for a skewed composition of amino acids, we explored two sequence recoding strategies on the alignment of the concatenated SCC proteins. In the first approach, we searched for a set of amino acid bins that “minimize the maximum chi-squared statistic” of heterogeneity, as implemented in the “MaxMin-ChiSq” program (Susko & Roger 2007). However, the program did not find any binning of amino acids that reduced heterogeneity in the alignment (homogeneity was rejected at p < 0.002 even for the highest-ranked bin choices), indicating that the alignments are still heterogenous despite the attempted recoding.

In the second approach, we manually selected a recoding that controls for IVYWREL bias in spirit of the Dayhoff 6-state recoding, which is commonly used to reduce heterogeneity while accounting for similarity in physiochemical properties of certain amino acids (Hrdy et al. 2004). The reconstructed phylogeny has the same major topological relationships at the genus and higher taxonomic levels in comparison to the tree obtained using non-recoded SCC proteins (**Supplementary Figure 6**).

Phylogenetic mixed models can be more effective at accounting for heterogeneity than recoding (Le et al. 2008; Hernandez & Ryan 2021). We used two different mixture models to test for changes in topologies: the Posterior Mean Site Frequency Model (PMSF) (Wang et al. 2018) and the General Heterogeneous evolution On a Single Topology (GHOST) model (Crotty et al. 2020), which could account for possible heterotachous evolution due to optimum growth temperature’s impact on mutation rates in different lineages and at different sites in a protein sequence (Zeldovich et al. 2007; Crotty et al. 2020). The topologies of trees reconstructed using both models (**Supplementary Figures 7** and **8**) are consistent with the SCC proteins’ tree built under a homogeneous model.

Overall, and despite the challenges of compositional biases, phylogenies of marker genes strongly support the early divergence of *Mesoaciditogales* from the last common ancestor of *Thermotogota*.

### Evolutionary histories of many *Mesoaciditogales* and *Petrotogales* genes are intertwined

The analyzed 232 single-copy gene families form only a small portion of any *Thermotogota* genome, which encode between 1,564 and 3,097 protein-coding genes across the described *Thermotogota* species (*M. lauensis* and *A. saccharophila* genomes encode 2,054 and 2,004 protein-coding genes, respectively). The histories of these conserved core genes often do not reflect the complex evolution of prokaryotic genomes, which are substantially impacted by horizontal gene transfer (Philippe & Douady 2003; Gogarten & Townsend 2005; Zhaxybayeva et al. 2009; Arnold et al. 2022; Doolittle & Brunet 2016). Therefore, we expanded our analyses to 1,181 gene families detected in both *M. lauensis* and *A. saccharophila*.

To investigate placement of *Mesoaciditogales* within *Thermotogota* on the individual gene family trees, we developed an approach that we dubbed the “minimal bipartition” analysis. In this method, we represent each gene family phylogeny as a set of bipartitions. The expected basal position of *Mesoaciditogales* would produce a bipartition that separates *Mesoaciditogales* and the outgroup taxa from other *Thermotogota*. However, in a tree with a non-basal position of *Mesoaciditogales* this bipartition would not exist. Our algorithm first identifies the tree bipartitions that contain *M. lauensis* and *A. saccharophila* and the outgroup taxa, and then finds the smallest set of other *Thermotogota* taxa that are required to create each split. This “minimum bipartition” process can quantify the relationship of *Mesoaciditogales* to other *Thermotogota* without visual inspection of individual gene trees. Of the 1,181 gene families, 721 (including 190 of 232 SCC families) met the criteria for a minimum bipartitions analysis (see **Methods** section for details).

In 234 of the 721 gene families’ trees (32%), *Mesoaciditogales* branch basally in the *Thermotogota* phylum (**Figure 4**), although 3 of these gene families are not found in the order *Thermotogales*. In 203 out of 721 trees (28%), *Petrotogales* taxa join the minimum bipartition (“*Mesoaciditogales + Petrotogales”* in **Figure 4**). Only 8 of these 203 families are found solely in *Mesoaciditogales* and *Petrotogales* taxa, indicating that many genes that are present across the *Thermotogota* phylum have evolutionary history similar to the 16S rRNA gene phylogeny discussed above. In 223 of the 721 trees (31%), minimum bipartition includes a mixture of *Petrotogales* and *Thermotogales* (“mixed-history” in **Figure 4**), suggesting that evolutionary histories of these gene families likely involve multiple horizontal gene transfer events, which are common in *Thermotogota* (Zhaxybayeva et al. 2009; Pollo et al. 2015).

**Figure 4.**
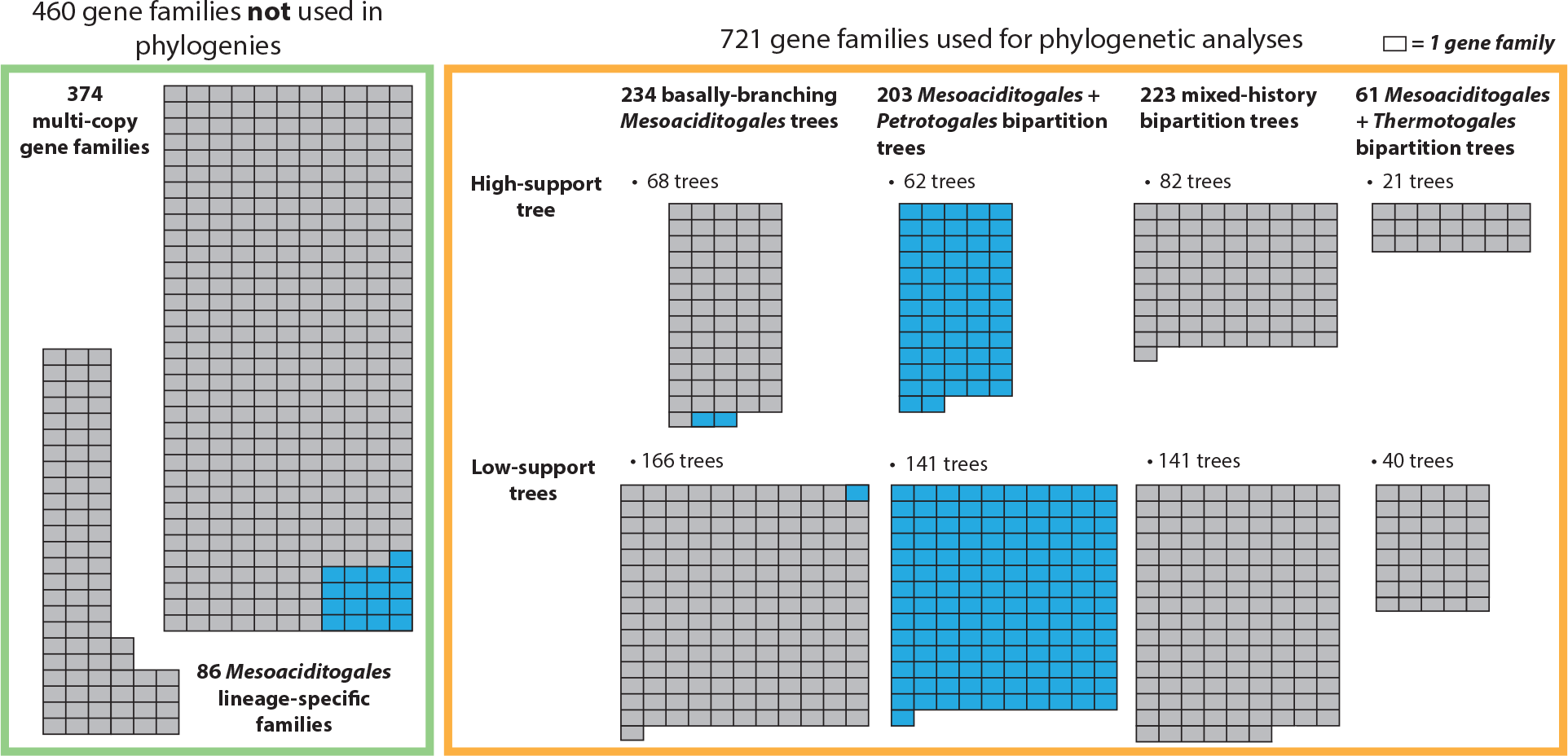
The distribution of phylogenetic relationships in *Thermotogota* gene families that contain *Mesoaciditogales*. Each rectangle represents one gene family. Gene families in the green box were analyzed only for taxonomy of their members. Gene families in orange box underwent phylogenetic reconstruction. The latter gene families are subdivided into categories based on the taxonomic composition of their “minimum bipartition” and the support of that bipartition (high support trees have QP- IC score ≥ 0.3 and bootstrap support ≥50%). *Petrotogales*-associated gene families (see text for definition) are in shown in blue.

Many of the 721 trees have low support for the above-described bipartitions, which may be due to short sequence length, limited number of informative sites, or poor alignment quality. However, 233 “high-support” trees (QP-IC score ≥ 0.3 and bootstrap support ≥50% for the minimum bipartition) result in similar fractions of trees supporting the above-described scenarios (**Figure 4**).

Additionally, among the 460 gene families not used for phylogenetic analyses, 17 are found exclusively in *Mesoaciditogales* and *Petrotogales* taxa (**Figure 4**). Combined with the 203 “*Mesoaciditogales + Petrotogales*” families and the 3 “basally-branching” gene families found exclusively in *Mesoaciditogales* and *Petrotogales* species, a total of 223 of 1,181 gene families found in both *Mesoaciditogales* genomes (19%) share their recent evolutionary history with their *Petrotogales* spp. homologs. We refer to this set of 223 families as “*Petrotogales*-associated” gene families.

### Genes that share recent evolutionary histories with *Petrotogales* have functions important for responding to environmental conditions

The 223 *Petrotogales*-associated gene families contain 243 *Mesoaciditoga lauensis* genes (**Supplementary File 1**), which can be further subdivided in two sets: 195 genes (from 195 gene families) that are present widely in the *Thermotogota* and 48 genes (from 28 gene families) found solely in *Mesoaciditogales* and *Petrotogales* taxa. Many of these 243 genes (as highlighted below) are known to be involved in response to environmental changes. We investigated functional connections among the 243 genes by reconstructing STRING association networks (Szklarczyk et al. 2015) and examining their Clusters of Orthologous Groups (COG) categories (Galperin et al. 2021).

In the set of 195 widely-distributed genes, products of 182 genes are predicted by STRING to interact with at least one other protein from the set (**Figure 5A**). The largest of these networks is enriched in genes involved in gene expression, translation, and protein export, and contains a highly connected subnetwork of genes relating to bacterial chemotaxis. In the set of 48 genes, 16 genes form a few small networks (**Figure 5B)**, including a network of genes involved in *de novo* UMP biosynthesis and nucleoside monophosphate and pyrimidine metabolism, several networks of genes involved in transport of amino acids and carbohydrates, a network of lipid metabolism genes, and a network related to toxin-antitoxin systems. Thus, many of the 243 *Petrotogales*-associated genes are likely functionally connected, or even belong to the same biochemical pathway or molecular complex.

**Figure 5.**
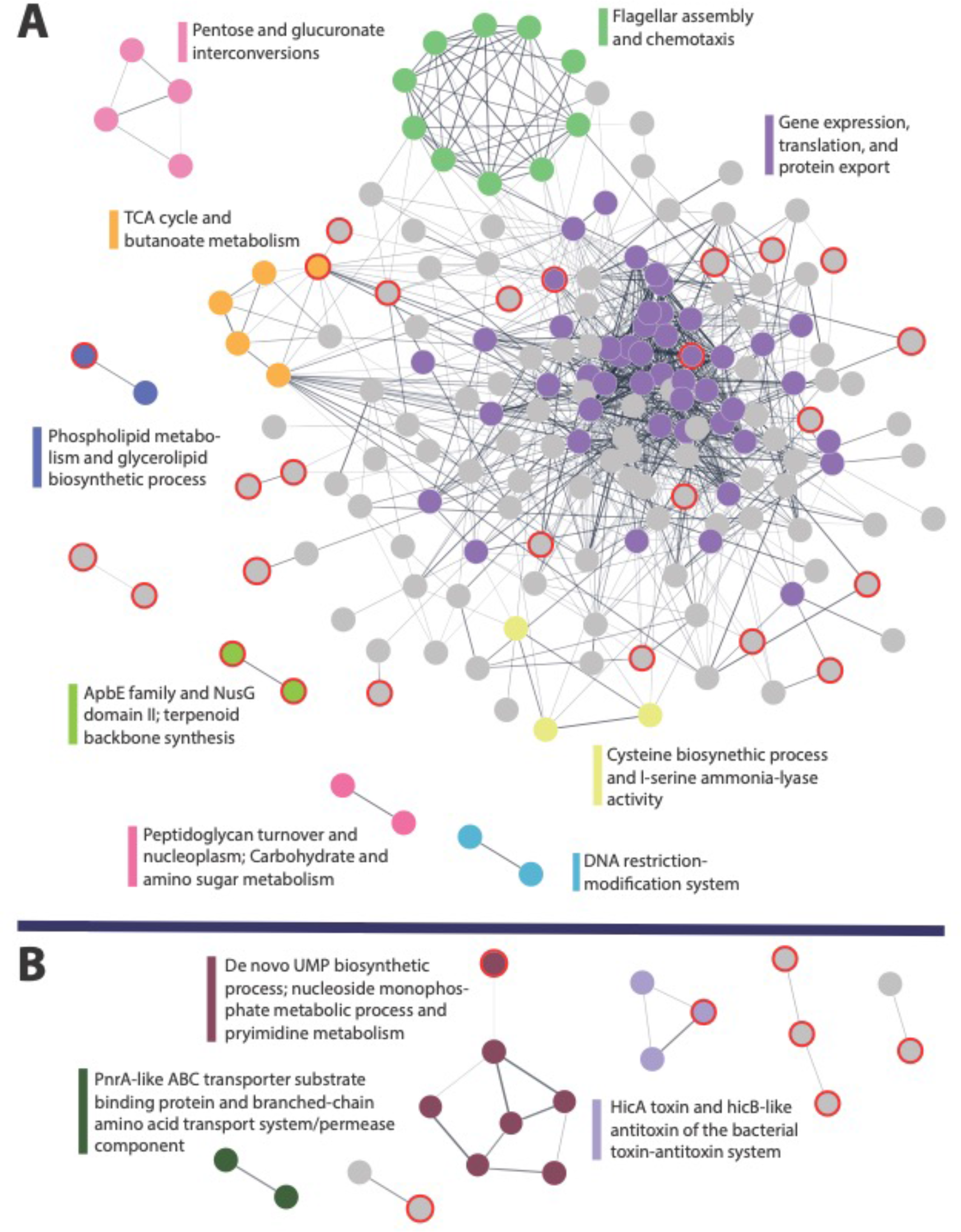
**The STRING protein-protein interaction networks of *M. lauensis* genes from the 223 *Petrotogales*-associated gene families**. **A**. Networks of genes that are present widely in *Thermotogota* (182 connected proteins). **B.** Networks of genes found solely in *Mesoaciditogales* and *Petrotogales* (22 connected proteins). Singleton proteins are not shown. The edges are weighted by the confidence of the interactions, with a minimum association threshold of 0.4. Subnetworks enriched for certain functions have nodes highlighted in color and their “STRING Cluster” annotations listed. Grey nodes are not part of enriched subnetworks. Nodes outlined in red are from proteins of unknown function.

The aforementioned genes for *de novo* UMP biosynthesis are assigned to the “coenzyme transport and metabolism” (H) COG category. Another gene in the H COG category, glyceraldehyde3-phosphate dehydrogenase, is known to be differentially regulated in *Kosmotoga olearia* (order *Petrotogales*) depending on temperature (Pollo et al. 2017). More broadly, genes from the H COG category are overrepresented among the 243 genes (Fisher’s exact test, p = 0.005). Within the STRING’s “gene expression, translation, and protein export” subnetwork and among 13 genes with post-translational modification, protein turnover, and chaperone functions (COG category O) are two genes which encode GroEL and a serine protease DO chaperone.

These two proteins are implicated in the response to heat stress in *K. olearia* (Pollo et al. 2017). Among genes assigned to “Translation, ribosomal structure and biogenesis” (J) COG category are genes encoding ribosomal protein L12P and ribosome binding factor A (*rbfA*), which, along with other ribosomal proteins, are known to exhibit changes in expression linked to cold response (Pollo et al. 2017; Jones & Inouye 1996; Barria et al. 2013). Two genes that form the small “peptidoglycan turnover and nucleoplasm” subnetwork (**Figure 5A)** are part of fatty acid synthesis and membrane envelope biogenesis (COG category M), along with 6 additional genes. One poorly characterized gene (COG category R), putatively annotated to encode enoyl-ACP reductase II, is connected to two lipid transport and metabolism genes (COG category I) within the large STRING network of *Petrotogales*-associated genes. In the *Mesoaciditogales* genomes, this gene is located in a neighborhood of genes involved in fatty acid biosynthesis, and is therefore likely involved in that biochemical pathway. Based on a gene neighborhood analysis using DOE IMG/M website (IMG Gene ID 2582540014; Accessed June 14 2023)(Chen et al. 2017), the gene is likely acquired from a Firmicute (*Thermincola*).

There are 55 additional genes of yet unknown function or with only general function assigned *in silico* (COG category R or S) (**Supplemental File 1**). Eleven of these genes have temperature-responsive homologs in *Kosmotoga olearia* (Pollo et al. 2017), and 26 are *Mesoaciditogales*-*Petrotogales* specific. Products of some of these uncharacterized genes are predicted to interact with other *Petrotogales*-associated genes (**Figure 5**). Hence, there are likely undiscovered genes important for specific environmental adaptations.

## Discussion

Our phylogenetic analyses of commonly used phylogenetic markers (16S rRNA and single-copy core genes) support the placement of the order *Mesoaciditogales* at the base of the *Thermotogota* phylum (**Figure 1** and **Supplementary Figure 1**). This inference does not change when taxa are added to break long branches, sequences are re-coded to adjust for compositional biases, and non-homogeneous models are used to correct for compositional heterogeneity across *Thermotogota* phylum.

However, expanding the analyses beyond these commonly used marker genes reveals that many gene families in *Mesoaciditogales* have alternative evolutionary histories (**Figure 4** and **6)**. It is not surprising that genomic content of *Mesoaciditogales* has been impacted by extensive HGT, as it is common in other *Thermotogota* (Zhaxybayeva et al. 2009; Nesbø et al. 2015; Haverkamp et al. 2021) as well as in bacteria in general (Philippe & Douady 2003; Gogarten & Townsend 2005; Arnold et al. 2022; Doolittle & Brunet 2016).

**Figure 6.**
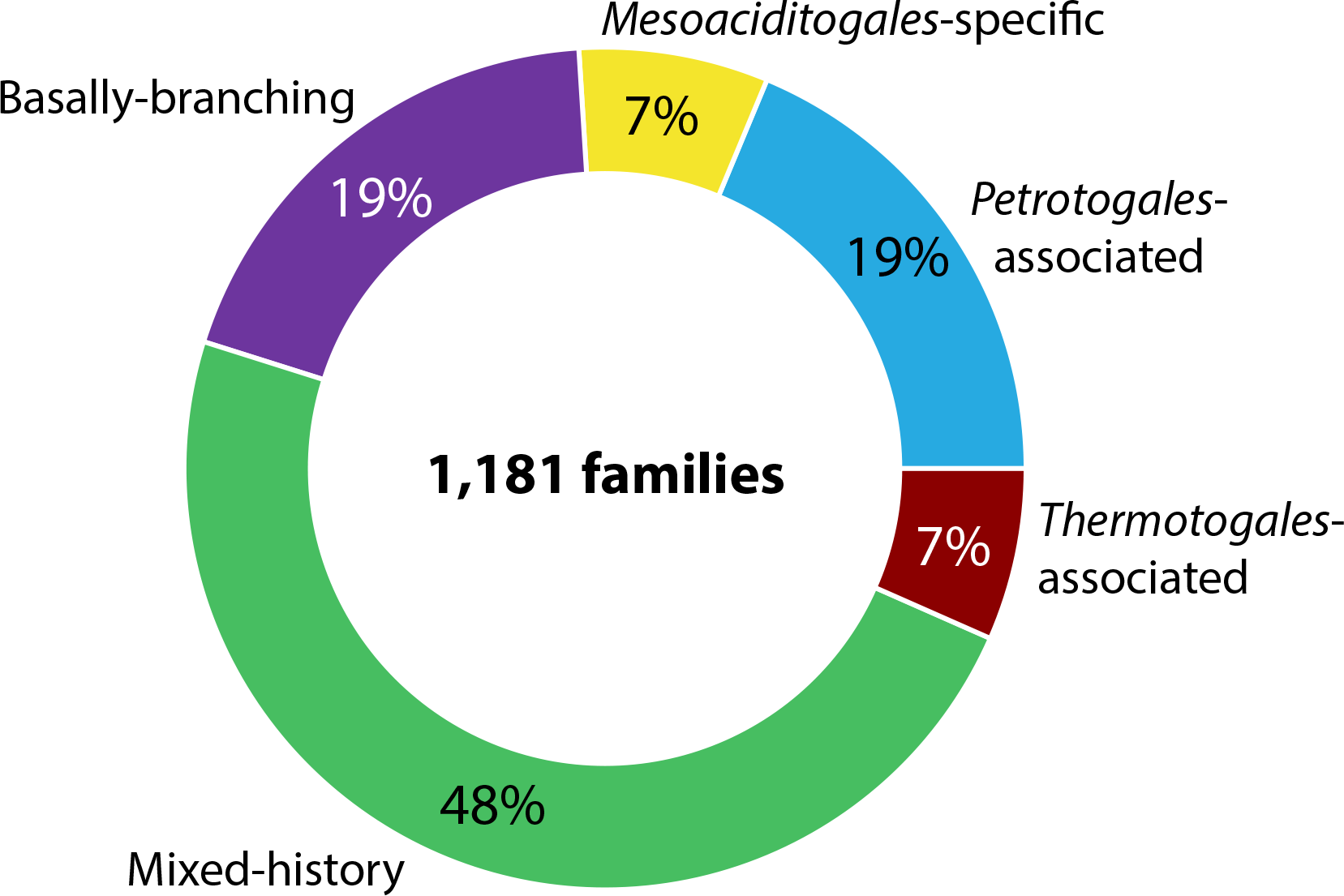
Summary of phylogenetic relationships of 1,181 gene families found in both *Mesoaciditogales* with respect to other *Thermotogota*.

Notably, nearly a fifth of *Mesoaciditogales* gene families are more closely related to taxa from *Petrotogales* order (**Figure 6**), many of which grow optimally at lower temperatures than members of *Thermotogales* order (**Figure 1)**. Some *Petrotogales* species have been found in similar environments as *Mesoaciditogales* and *Mesoaciditogales*-like MAGs (L’Haridon et al. 2019; Mori et al. 2020; Nesbø et al. 2021; L’Haridon et al. 2023), which could provide an opportunity for HGT between members of these two taxonomic groups. Alternatively, the *Mesoaciditogales* and *Petrotogales* lineages may have independently acquired these homologs via other taxa, such as *Firmicutes*.

We hypothesize that evolutionary histories of these genes reflect HGT-facilitated adaptation of *Kosmotogales* and *Mesoaciditogales* to similar ecological niches. For instance, *Mesoaciditoga lauensis* and *Kosmotogaceae* species share genes needed to grow on acetate that are not found in other *Thermotogota* species (Nesbø et al. 2019). In another example, *de novo* UMP biosynthesis is a highly conserved process that requires specific adaptations in hyperthermophilic environments (Thia-Toong et al. 2002) – which may explain why homologs of *Mesoaciditogales* UMP-related genes are found in *Petrotogales*, but not in the hyperthermophilic *Thermotogales*. Other central carbon metabolism genes, such as glyceraldehyde3-phosphate dehydrogenase and the small networks of metabolism-related genes, may be specifically adapted to “ideal” environmental conditions, since metabolism is maximally upregulated at optimal temperatures and growth conditions (Pollo et al. 2017). The similarity of ribosome-associated genes (encoding ribosomal proteins and ribosome binding factor A) suggests that the ribosomes in *Mesoaciditogales* and *Petrotogales* might be fine-tuned to their preferred growth temperature range, consistent with earlier studies of temperature adaptations of *Kosmotoga olearia* and other bacteria (Barria et al. 2013; Pollo et al. 2017).

Several genes that are known to have temperature-dependent expression in *Kosmotoga olearia* were among the *Petrotogales*-associated genes. Glyceraldehyde3-phosphate dehydrogenase is upregulated in *Kosmotoga olearia* (order *Petrotogales*) at its OGT of 65°C (Pollo et al. 2017) – a temperature close to *Mesoaciditoga*’s OGT. The two genes that are known to be part of *Kosmotoga olearia*’s heat stress response, GroEL and a serine protease DO chaperone, both gene have homologs across *Thermotogota*, but the similarity between these *Petrotogales* and *Mesoaciditogales* homologs may be due to specialized adaptations to specific ranges of temperature stress, or they may have shared their evolutionary history through HGT. In *Kosmotoga olearia,* enoyl-ACP reductase II contributes to changes in the cellular membrane composition in response to growth temperature, and its gene is also predicted to be horizontally acquired (Pollo et al. 2017). This and other *Mesoaciditogales*’ and *Petrotogales-*associated genes related to fatty acid synthesis, membrane biogenesis, and lipid metabolism are likely important for membrane remodeling in response to shifting temperatures (Suutari & Laakso 1994; Pollo et al. 2017). While these genes represent only a fraction of the 243 *Petrotogales*-associated genes, many of the remaining genes are predicted to have functional associations with the highlighted genes. This raises a possibility that their involvement in environmental adaptations may have not yet been identified.

Since HGT likely had a strong role in shaping *Mesoaciditogales’* genome content, the question about the OGT of the last common ancestor of *Thermotogota* remains open. It is possible that within *Thermotogota* adaptations to lower temperature happened multiple times, independently, and that the LCA of the phylum is a hyperthermophile as previously hypothesized (Zhaxybayeva et al. 2009; Green et al. 2013). To resolve this question, further analyses of gene gain, loss and HGT across the phylum are needed.

## Methods

### Retrieval and analyses of 16S rRNA genes

*Thermotogota* 16S rRNA gene sequences from 55 described *Thermotogota* species, whose strains are available in the DSMZ collection [https://www.dsmz.de/], were downloaded from GenBank (Benson et al. 2018) on November 8, 2020. Eight sequences were added as an outgroup: *Hydrogenivirga caldilitoris* IBSK3 (NR_024824), *Aquifex aeolicus* VF5 (NR_114796), *Hydrogenobacter thermophilus* TK-6 (NR_074870), *Thermocrinis ruber* OC 1/4 (NR_121741), *Persephonella marina* EX-H1 (NR_102828), *Dictyoglomus thermophilum* H-6-12 (NR_029235), *Staphylococcus aureus* ATCC 12600 (NR_118997) and *Coprothermobacter platensis* (Y08935). The sequences were aligned using SILVA’s Incremental Aligner (SINA) v1.2.11 (Pruesse, Peplies and Glöckner, 2012). The multiple sequence alignment was trimmed using trimAl v1.4.rev22 with the –gappy-out setting (Capella-Gutiérrez et al. 2009), resulting in the 1,548 nt alignment. Pairwise nucleotide identity among the *Thermotogota* 16S rRNA genes was calculated from the trimmed alignment using in-house scripts.

The dataset was expanded by adding environmental sequences, which were retrieved using *Mesoaciditoga lauensis* cd-1655R^T^ 16S rRNA gene (NR_125611) as a query in a web-based BLASTN search (Altschul et al. 1990) of the *nt* database (Benson et al. 2018) with low complexity filter on, E-value threshold of 0.0001, a maximum of 5,000 target sequences, and all other parameters as default (the search was performed in February 2021). All matches with >80% sequence identity to the query were added to the dataset of 16S rRNA genes from 55 described *Thermotogota* species and the whole dataset was re-aligned with SINA v1.2.11 (Pruesse, Peplies and Glöckner, 2012). This resulted in an alignment of 277 sequences that contains 2,168 sites.

The maximum likelihood (ML) tree under a homogeneous model was constructed in IQ-TREE v1.6.7 (Nguyen et al. 2015) under the GTR+F+R4 model chosen by ModelFinder (Kalyaanamoorthy et al. 2017). Accuracy of the topology was assessed via bootstrap analysis of 100 pseudo-samples.

The heterogeneity of sequence composition was assessed using two methodologies. First, chi-square test was performed on individual sequences, as implemented in IQ-TREE v1.6.7 (Nguyen et al. 2015). Second, Bowker’s Test for non-stationarity was performed on whole dataset using TestNH v.1.3.0 from the Bio++ suite (Dutheil & Boussau 2008). The ML tree and model (GTR+G4 substitution model, alpha = 0.6) reconstructed in IQ-TREE (see above) was used as the starting tree, and a homogenous model was fit using the ML method implemented in the Bio++ bppml program. Then, the Bowker’s test was performed on the Bio++ homogenous tree with a p-value threshold of 0.05 using 1,000 parametric bootstraps.

A ML tree reconstruction under a non-homogenous model was performed using nhPhyML v0.1 (Boussau & Gouy 2006). Transition/transversion ratio was estimated from the dataset and among site rate variation was modeled under the G4 model. The rooted tree reconstructed under the homogenous model in IQ-TREE was provided as the starting tree.

Stem regions of the 16S rRNA genes were predicted using Infernal v.1.1.4 (Nawrocki & Eddy 2013). GC-content of full-length 16S rRNA genes and of their stem regions were calculated using in-house scripts.

### Retrieval of genomes and assignment of taxonomy

*Thermotogota* genomes and metagenome-assembled genomes (MAGs) (412 in total) were obtained either from NCBI’s “Assembly” database (Kitts et al. 2016) or from the IMG database (Chen et al. 2017) in December 2020. Duplicated genomes found in both databases were removed. Additionally, 42 MAGs were shared with us privately by Eric Boyd (Montana State University), Håkon Dahle (University of Bergen), and Brett Baker (University of Texas in Austin). The genome list was checked against the list of strains of *Thermotogota* available in the DSMZ collection (https://www.dsmz.de/, last accessed December 2020) to ensure the inclusion of all described species with available genomes, and against the NCBI’s Taxonomy database (Schoch et al. 2020) to verify that MAGs were classified as *Thermotogota* (NCBI taxid 200918). Genomes were assessed for completeness using CheckM (Parks et al. 2015), and only genomes that were estimated to be at least 50% complete were retained. Fifteen genomes from bacteria belonging to other phyla were used as an outgroup. Since *Thermotogota* are strictly anaerobic bacteria, the aerobic taxa were included in the outgroup to reduce the potential impact of HGT on gene phylogenies.

This procedure resulted in 172 genomes (154 genomes and MAGs of *Thermotogota,* 15 genomes of outgroup species, and 3 unassigned MAGs that grouped with the outgroup), which were used in the subsequent analyses. GenBank and IMG accession numbers, as well as genome completeness information, are listed in **Supplementary File 2**.

Genomes were assigned taxonomy based on the Genome Taxonomy Database (GTDB)(Chaumeil et al. 2020). While NCBI’s Taxonomy database (Schoch et al. 2020) classifies *Kosmotogaceae* as a family within the order *Kosmotogales*, the GTDB nomenclature places both the *Kosmotogaceae* and *Petrotogaceae* families into the order *Petrotogales.* We use the GTDB nomenclature throughout this manuscript. Sixty of the 172 genomes belong to strains of described *Thermotogota* species.

### Phylogenetic reconstruction from genes encoding ribosomal proteins

Amino acid sequences of genes encoding ribosomal proteins were identified by using 50 *Thermotoga maritima* ribosomal proteins as queries in BLASTP v. 2.6.0 searches (Altschul et al. 1990) against the dataset of the above-described 172 genomes. Results were filtered for coverage and quality (e-value < 0.0001, and the match length is between 60 and 140% of the query length), and then verified to be annotated as ribosomal proteins.

For the 60 strains of described *Thermotogota* species and all 15 outgroup species, the retrieved sequences for each of the 50 ribosomal proteins were aligned in MAFFT v7.305b using-linsi setting (Katoh & Standley 2013). The alignments were concatenated with in-house scripts into one alignment; in cases where a genome did not have a detected ribosomal protein, gaps were inserted into the alignment. The phylogenetic tree was reconstructed using the trimmed concatenated alignment in IQ-TREE v1.6.7 (Nguyen et al. 2015) using the LG+F+R5 substitution model selected by built-in ModelFinder (Kalyaanamoorthy et al. 2017). Accuracy of the topology was assessed via bootstrap analysis of 100 pseudo-samples.

The alignment, concatenation, and tree reconstruction were repeated using the 50 ribosomal proteins from all 172 genomes to assess how the inclusion of MAGs would affect the topology. For this expanded phylogenetic analysis, the LG+F+R5 model was again selected by ModelFinder (Kalyaanamoorthy et al. 2017).

### Likelihood mapping

Likelihood mapping analyses for 16S rRNA gene and ribosomal protein alignments were carried out in TREE-PUZZLE 5.3.rc16 (Schmidt & Haeseler 2007).

For 16S rRNA sequence analyses, the trimmed alignment of sequences from described species was used (1,548 nucleotide sites for 63 rRNA sequences, including outgroup). All 11,232 possible quartets were evaluated using approximate parameter estimation for computational efficiency. GTR model was used for the substitution model with nucleotide frequencies estimated from the alignment, and rate heterogeneity modeled using Gamma distribution with four rate categories. Two tests were performed: In the first test, all sequences were assigned into one of the four taxonomic groups, *Mesoaciditogales*, *Thermotogales*, *Petrotogales*, or the outgroup species; in the second test, sequences from *Thermotogota* were divided into *Mesoaciditogales*, *Thermotogales*, *Petrotogaceae,* or *Kosmotogaceae,* while the outgroup species were excluded.

For the ribosomal proteins analyses, all 50 protein alignments were analyzed individually. For each protein, all possible quartets were analyzed using approximate parameter estimation for computational efficiency. The LG substitution matrix was used for the substitution model with amino acid frequencies estimated from the alignment, and rate heterogeneity modeled using Gamma distribution with four rate categories. All sequences were assigned to one of four taxonomic groups: *Mesoaciditogales*, *Thermotogales*, *Petrotogales*, or outgroup.

### Calculation of bias towards IVYWREL amino acids

For each protein encoded in 60 genomes from described *Thermotogota* species, the proportion of IVYWREL amino acids was calculated using in-house scripts (available in the **FigShare repository**). Transmembrane proteins were predicted using Phobius v.1.01 (Käll et al. 2004). The linear regression analysis of median IVYWREL of the non-transmembrane proteins and optimum growth temperature of the 60 species was performed using scikit-learn v.0.23.2 (Pedregosa et al. 2011). Optimal growth temperatures were retrieved from each species’ defining publication and cross-checked against strain and species data from BacDive (Reimer et al. 2019) when available.

### Identification of Gene Families

Gene families (orthogroups) in the 172 genome dataset were identified in OrthoFinder v.2.5.1 (Emms & Kelly 2019). Within OrthoFinder, BLASTP was used for the sequence search, and MAFFT v.7.305b for sequence alignment. FASTA files for gene families are available in the **FigShare repository**.

### Analyses of single-copy gene families

Single-copy gene families conserved across the *Thermotogota* phylum (“Single-copy core”, or SCC, gene families) were defined as orthogroups that contained genes from at least 45 of the 60 genomes from described *Thermotogota* species. This threshold was chosen to maximize the number of gene families, while retaining genes that are found across the majority (>50%) of taxa in each of the three *Thermotogota* orders (**Figure 1**). The selection resulted in 232 gene families, which include all 50 ribosomal proteins described above. The sequence alignments were retrieved from OrthoFinder and were concatenated using in-house scripts. The concatenated alignment was trimmed with TrimAl v1.4.rev22 using the -gappyout setting (Capella-Gutiérrez et al. 2009). This trimmed concatenated alignment was used throughout the phylogenetic analyses described in this section.

Compositional heterogeneity of sequences was assessed using the Chi-squared testing performed in IQ-TREE v1.6.12 (Nguyen et al. 2015). The phylogenetic tree was reconstructed in IQ-TREE v1.6.12 using the LG+F+R8 model selected by ModelFinder (Kalyaanamoorthy et al. 2017). Accuracy of the topology was assessed via bootstrap analysis of 100 pseudo-samples.

Alternative topology testing was carried out in IQ-TREE v1.6.12 (Nguyen et al. 2015). The SCC protein alignment was used to reconstruct maximum likelihood phylogenetic trees that followed a set of topological constraints, which were imposed to define various relationships among *Thermotogota* families (**Figure 2**). All trees were reconstructed under the LG+F+R8 model, which was selected as the best fitting model for the SCC alignment during the unconstrained tree reconstruction. The likelihoods of the constrained trees were compared to the likelihood of the unconstrained SCC proteins phylogeny using the approximately unbiased (AU) test (Shimodaira 2002).

The alignment recoding was carried out using two approaches. In the first approach, recoding states that minimize sequence heterogeneity were searched using MinMax-ChiSq v1.1 (Susko & Roger 2007), which assessed all bin sizes between 2 and 20 amino acids and considered 5,000 random choices of starting bins for each bin size. In the second approach, recoding was used to adjust for the proportion of IVYWREL amino acids. Specifically, amino acids were recoded into 10 states (A,G,P,S,T,C, D**E**NQ, HK**R**, **IL**M**V**, F**WY**). This 10-state recoding is modified from the Dayhoff recodings (Hrdy et al. 2004; Embley 2003), which groups amino acids into states based on physiochemical characteristics. Per recommendation by (Hernandez & Ryan 2021), the model was tailored to our specific case as follows. Four states that involve IVYWREL amino acids as defined by Dayhoff 6-state recoding (D**E**NQ, HK**R**, **IL**M**V**, F**WY**) were retained, since IVYWREL bias is correlated with OGT in *Thermotogota.* The remaining 6 amino acids were allowed to be their own states since frequencies of these amino acids in proteins are presumed to be less affected by OGT. The recoding was carried out using in-house scripts (available in the **FigShare repository**). The phylogeny was built with RAxML v8.2.11 using the GTR+G model for multistate inference.

Phylogenetic trees were also reconstructed under two non-homogenous phylogenetic models as implemented in IQ-TREE v1.6.12 (Nguyen et al. 2015). First, the posterior mean site frequency (PMSF) model was used to account for site-specific compositional frequencies. The SCC phylogeny was provided as a guide tree to compute the PMSF amino acid profiles for each alignment site. A phylogeny was built using the obtained profiles under the LG+C20+F+G model. Second, the heterogeneous, edge-unlinked General Heterogeneous evolution On a Single Topology (GHOST) model (Crotty et al. 2020), was used to correct for possible heterotachy.

Specifically, a phylogeny was built under the LG+F0*H4 model. The LG substitution model was selected due to it being the best fit substitution matrix under the homogeneous model; F0 was used to assign separate base frequencies to each class, and *H4 was used to estimate model parameters separately for each of the 4 mixture classes. Accuracy of the reconstructions under both models was assessed using 1,000 ultrafast bootstrapping replicates.

### Identification of minimum bipartitions

For these analyses, 721 gene families that contain a single gene from *M. lauensis* genome, a single gene from *Athalassotoga saccharophila* genome, sequences from >5 additional *Thermotogota* genomes, and at least 3 outgroup sequences were selected. For each gene family, the maximum likelihood tree was reconstructed in IQ-TREE v. v1.6.12 using the MAFFT sequence alignment calculated in OrthoFinder and the best fitting substitution model selected by ModelFinder (Kalyaanamoorthy et al. 2017). Accuracy of the tree reconstruction was determined by analysis of 100 bootstrap replicates. Additionally, a quartet-based internode certainty score was used to complement bootstrap support by providing a measure of branch incongruence alongside the measure of topological accuracy (Zhou et al. 2020). This “QuadriPartition Internode Certainty” (QP-IC) was calculated using the maximum likelihood tree and the trees from bootstrap replicates.

Phylogenetic trees were converted to a set of bipartitions using bitstring representation in DendroPy v.4.5.2 (Sukumaran & Holder 2010). First, for each gene family tree, the largest bipartition that contained only outgroup taxa was labeled as the “outgroup bipartition”. Gene families with outgroup bipartition that contained less than 3 taxa were excluded from further analyses. Then, all bipartitions that included *M. lauensis, A. saccharophila,* and the taxa in the outgroup bipartitions were identified. Among these bipartitions, the bipartition that contained the smallest number of other taxa was labeled as the “minimum bipartition”. Trees of 20 gene families, in which *M. lauensis* and *A. saccharophila* did not group together, were excluded from further analyses. For each minimum bipartition, we identified the lowest taxonomic rank of the set of *Thermotogota* taxa that joined the *Mesoaciditogales.* Taxonomic rank of the set was determined by using the GTDB taxonomy classification of the 60 strains from described species.

Bipartitions were considered “highly-supported” if their QP-IC score ≥ 0.3 and bootstrap support ≥50%, and otherwise were considered “low-support” bipartitions. The analyses were carried out using in-house scripts (available in the **FigShare repository**).

### Identification of taxonomic rank for gene family composition

We assigned a lowest taxonomic rank to the 1,181 gene families containing *Mesoaciditogales*, including gene families not used in the phylogenetic analyses. Taxonomic rank was determined by excluding all *Mesoaciditogales* genomes, and then using GTDB taxonomic classification in conjunction with ETE 3 (Huerta-Cepas et al. 2016) to summarize which *Thermotogota* genera outside the *Mesoaciditogales* contributed genes to each family.

### COG assignment and COG categories’ enrichment calculations

Protein-coding genes from *M. lauensis* and *A. saccharophila* genomes were used as queries in the BLASTP v.2.6.0+ search of the NCBI COG database (2020 release; downloaded in March 2021)(Galperin et al. 2021). Forty-one percent of genes in these two genomes received a COG category assignment.

The distribution of COG categories in 1,181 gene families that contained both *Mesoacidotogales* species was compared to the distribution of COG categories in a subset of 223 “*Petrotogales*-associated” gene families (see **Results** for definition). Families were assigned a COG category by using the COG category of the *Mesoaciditogales* gene as a proxy. The significance of the difference was assessed using a one-sided Fisher’s Exact test, with a p-value threshold of ≤ 0.05.

### Reconstruction of protein-protein association networks

Protein-protein association networks were inferred using the STRING database and its web-based tools (Szklarczyk et al. 2015) (accessed on July 21, 2022). At the time of the analyses, the STRING database included the genome of *Mesoaciditoga lauensis* cd-1655R but did not contain the genome of *Athalassotoga saccharophila* genome. Therefore, the networks were reconstructed using only *M. lauensis* protein-coding genes. Specifically, functional associations were searched within the set of the 243 *M. lauensis* genes encompassed in the 223 “*Petrotogales*-associated” gene families (**Supplementary File 1**).

The 243 genes were separated into two sets: genes from families that also included genes from *Thermotogales* (195 genes), and genes from families that were present only in *Mesoaciditogales* and *Petrotogales* (48 genes). For each set, the separate association networks were inferred. For both networks, the default association threshold was used (0.4). Gene interaction tables, functional annotation files, and images of the original STRING networks are available in the **FigShare** respository. The two STRING association networks were imported into Cytoscape v.3.8 (Su et al. 2014), and the Cytoscape tool “StringApp” (Doncheva et al. 2019) was used to retrieve functional annotations of selected clusters within each network. The functions for all genes in *Mesoaciditogal lauensis’* genome were used as the background when calculating enrichment values.

### Phylogenetic Tree Visualization

All trees were visualized with iTOL’s web platform version 6 (Letunic & Bork 2021).

### Data Availability

The genomes and 16S rRNA sequences used for this research are publicly available via either NCBI Nucleotide (https://www.ncbi.nlm.nih.gov/nucleotide/), NCBI Assembly (https://www.ncbi.nlm.nih.gov/assembly) or the Joint Genome Institute Integrated Microbial Genomes (https://img.jgi.doe.gov/). Out of the 172 non-redundant *Thermotogota* genomes and MAGs used in this study, 141 are publicly available and their accession numbers are listed in **Supplementary File 2**. For the remaining 31 MAGs shared with us but not yet publicly available, the genes used in our analyses are available as part of the orthogroup data, described below. Amino acid sequences of proteins in orthogroups and 16S rRNA gene sequences in FASTA format, alignments, phylogenetic trees in Newick format, likelihood mapping results, STRING network analyses files, program log files, and in-house scripts are deposited in a **FigShare repository** available at DOI: 10.6084/m9.figshare.23303717.

## Acknowledgements

We thank Eric Boyd (Montana State University), Håkon Dahle (University of Bergen), and Brett Baker (University of Texas in Austin) for sharing MAGs, from which we extracted *Mesoaciditogales*’ homologs used in this study. The work was supported by Dartmouth Fellowship and Cramer funds to AAF, and Dartmouth start-up funds to OZ.

