## Supplementary Figures and Tables for "Early divergence and gene exchange highways in the evolutionary history of *Mesoaciditogales*"

**for**

**by**

Anne A. Farrell[1], Camilla L. Nesbø[2,3], Olga Zhaxybayeva[1,4,*]

1. Department of Biological Sciences, Dartmouth College, Hanover NH, USA.

2. Department of Biological Sciences, University of Alberta, Alberta, Canada.

3. Department of Chemical Engineering and Applied Chemistry, University of Toronto, Ontario, ON, Canada.

4. Department of Computer Science, Dartmouth College, Hanover NH, USA.

### Supplementary Figure 1 (panel A)

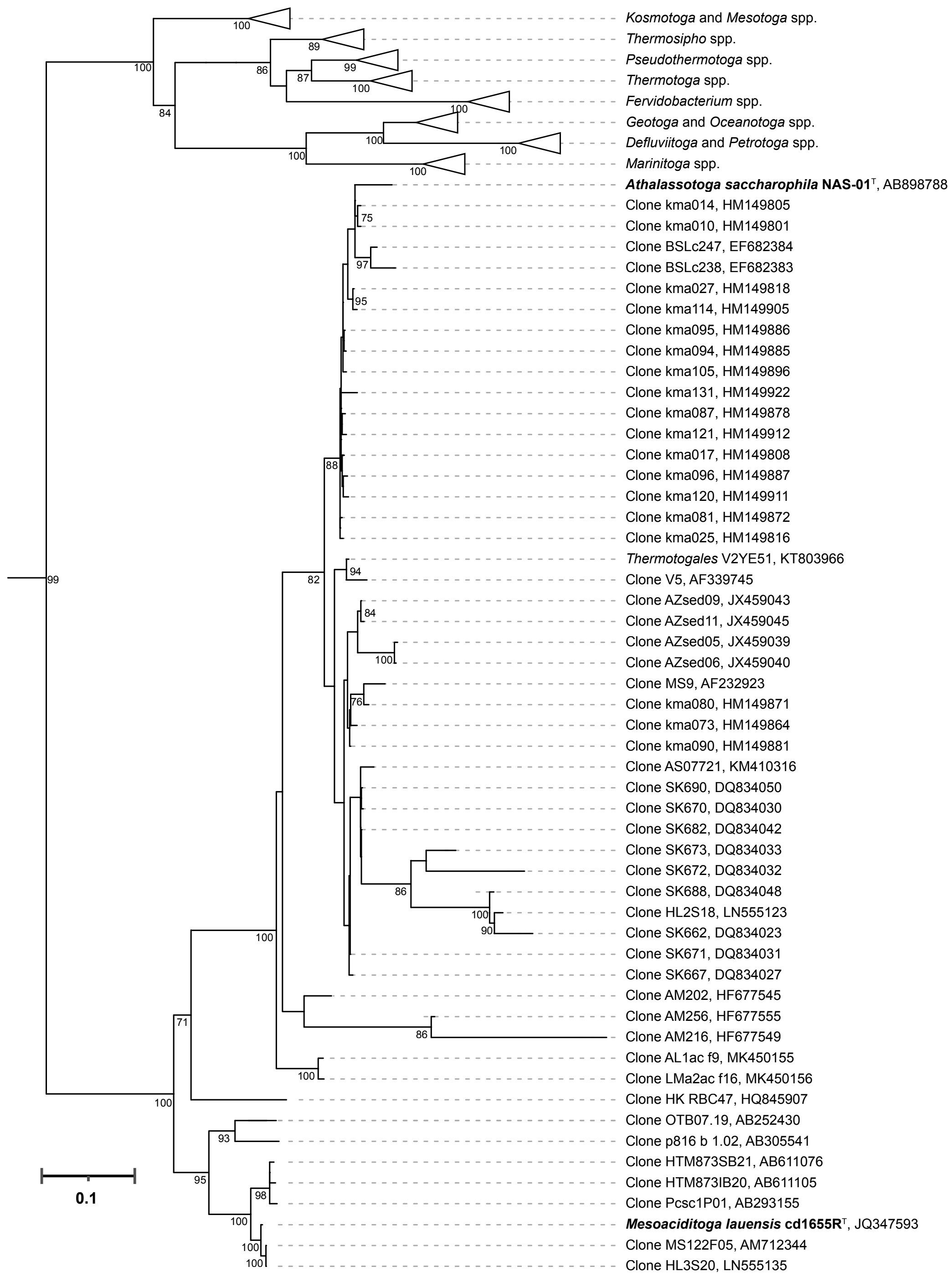

### Supplementary Figure 1 (panel B)

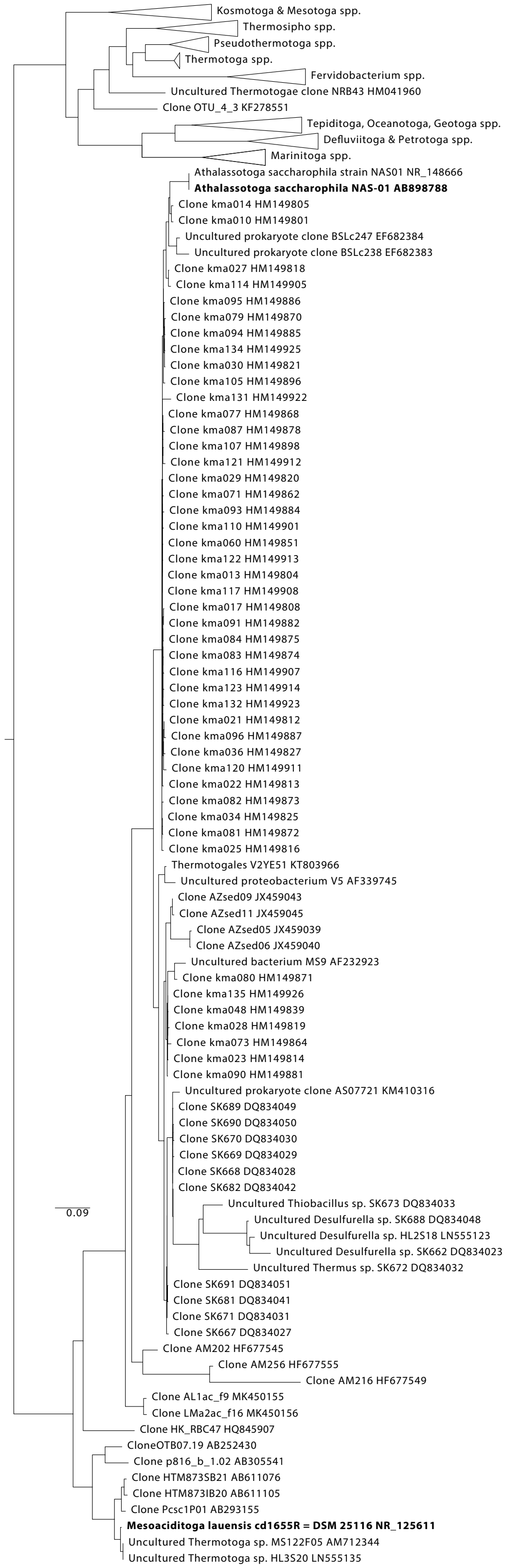

**Supplementary Figure 1. 16S rRNA gene phylogenies of described *Thermotogota* species, expanded to include environmental sequences closely related to *Mesoaciditogales*. (A)** The maximum likelihood tree reconstructed under the homogeneous (GTR+Γ+R4) substitution model. Bootstrap support values above 70% are shown. To avoid redundancy, clades of taxa with >99% sequence identity and originating from the same location are reduced to a single representative taxon by manual trimming of taxa, and taxa outside of *Mesoaciditogales* are collapsed. *Mesoaciditogales*' type strains are shown in bold font. The tree is rooted using an outgroup (not shown). GenBank accession numbers are listed after the taxonomic name. **(B)** The maximum likelihood tree reconstructed under non-homogeneous substitution model. Taxa outside of *Mesoaciditogales* are collapsed, and the tree is rooted using an outgroup (not shown). Trees from both panels are available uncollapsed, in Newick format in the **FigShare repository** (see **Methods** for details). Scale, substitutions per site.

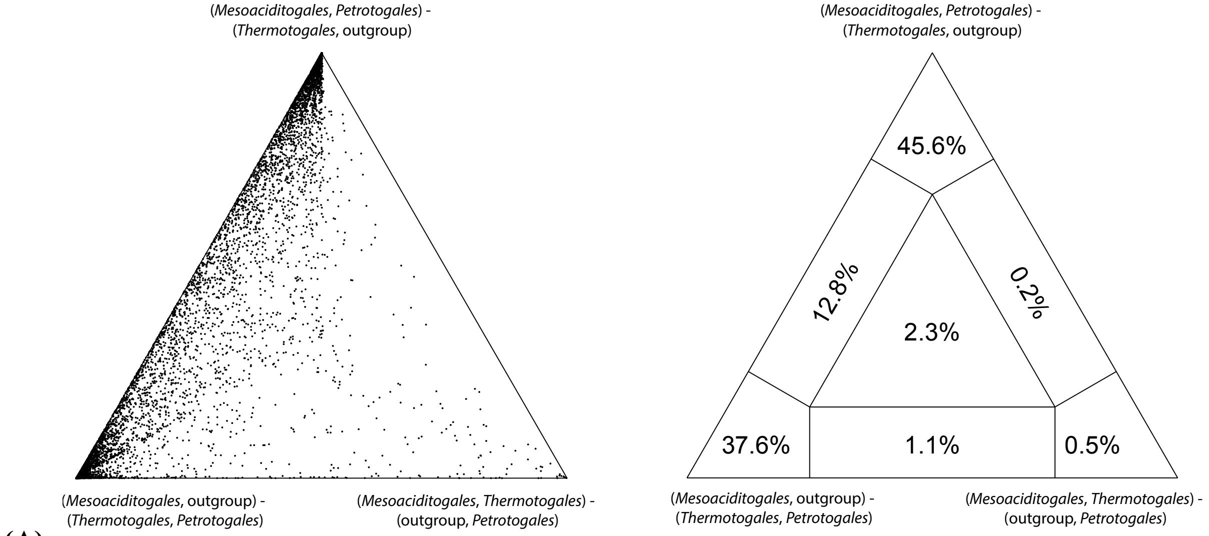


**(A)**

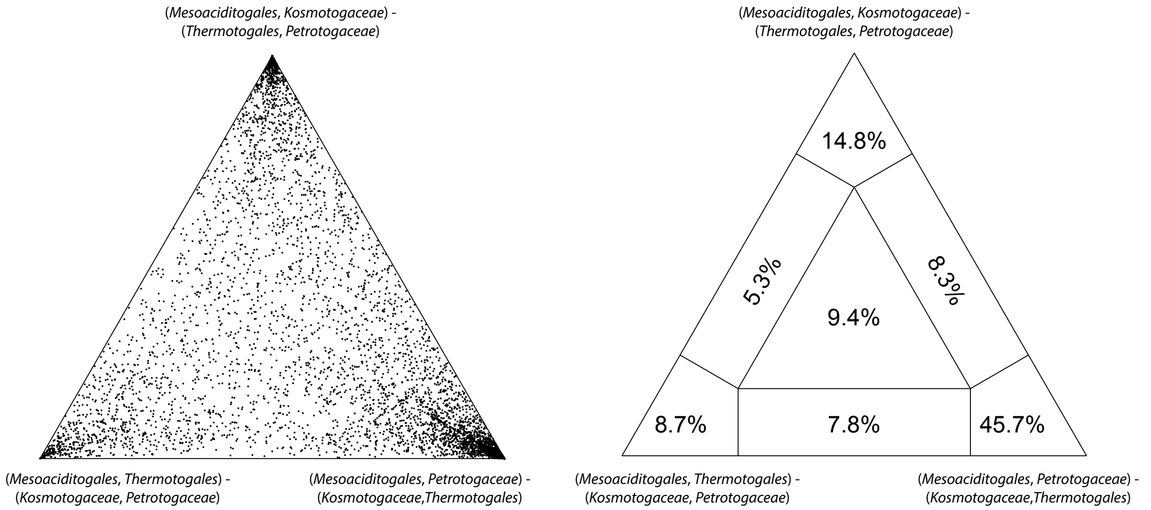


**(B)**

**Supplementary Figure 2. Likelihood mapping of 16S rRNA gene dataset of described *Thermotogota* species and the outgroup. (A)** Relationship among *Mesoaciditogales*, *Petrotogales*, *Thermotogales* and the outgroup. **(B)** Relationships among *Mesoaciditogales*, *Petrotogaceae, Kosmotogaceae,* and *Thermotogales* when the outgroup was excluded from the analyses. The three possible relationships among the tested groups are placed at the vertices of each simplex. In the simplexes on the left each dot represents a quartet of taxa in the dataset. The position of each dot in the simplex (barycentric coordinates) is determined by the vector of the support values for each of the three possible taxa relationships. The closer the dot to a vertex, the more support there is for the taxa relationship at the vertex. The simplex on the right summarizes the proportion of dots in each basin of attraction of the simplex, as defined by Strimmer and von Haeseler (1997).

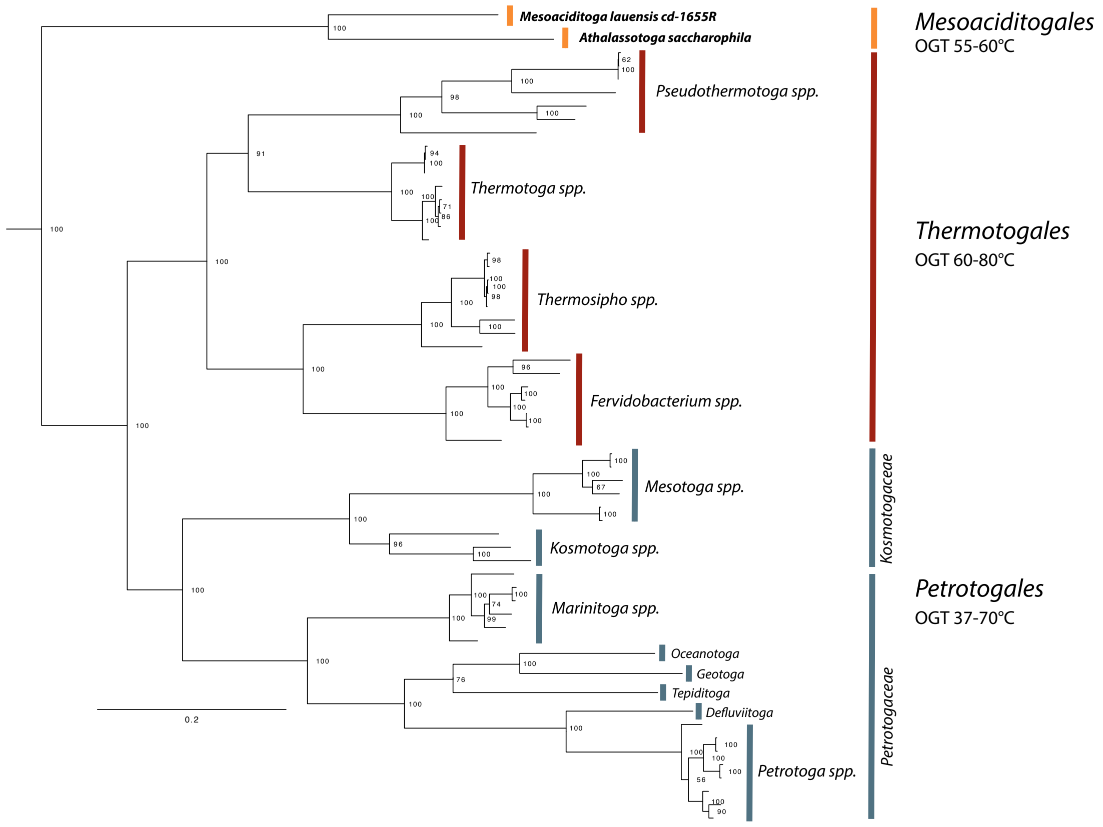


**Supplementary Figure 3. Concatenated ribosomal protein phylogeny for described *Thermotogota* species.** The maximum likelihood tree was reconstructed under the LG+F+R5 substitution model. *Thermotogales Petrotogales* and *Mesoaciditogales* genera are demarkated by red, blue, and orange bars, respectively. Numbers at the nodes represent bootstrap support values. The tree was rooted using outgroup taxa *Clostridium acetobutylicum* (GCA_000008765), *C. botulinum* (GCA_000063585), *Alkaliphilus metalliredigens* (GCA_000016985), *Bacillus velezensis* (GCA_000875875), *B. cereus* (GCA_003122725), *Saccharibacillus sacchari* (GCF_000585395), *Brevibacillus brevis* (GCF_003385915), *Pseudomonas fluorescens* (GCF_003701995), *Thermus thermophilus* (GCA_000091545), *Marinithermus hydrothermalis* (GCA_000195335), *Aquifex aeolicus* (GCF_000008625), *Persephonella marina* (GCF_000021565), *Dictyoglomus thermophilum* (GCF_000020965), and *D. turgidum* (GCF_000021645). The outgroup is not shown. Scale bar, substitutions per site. Tree with all taxa labels is available in Newick format in the **FigShare repository** (see **Methods** for details.)

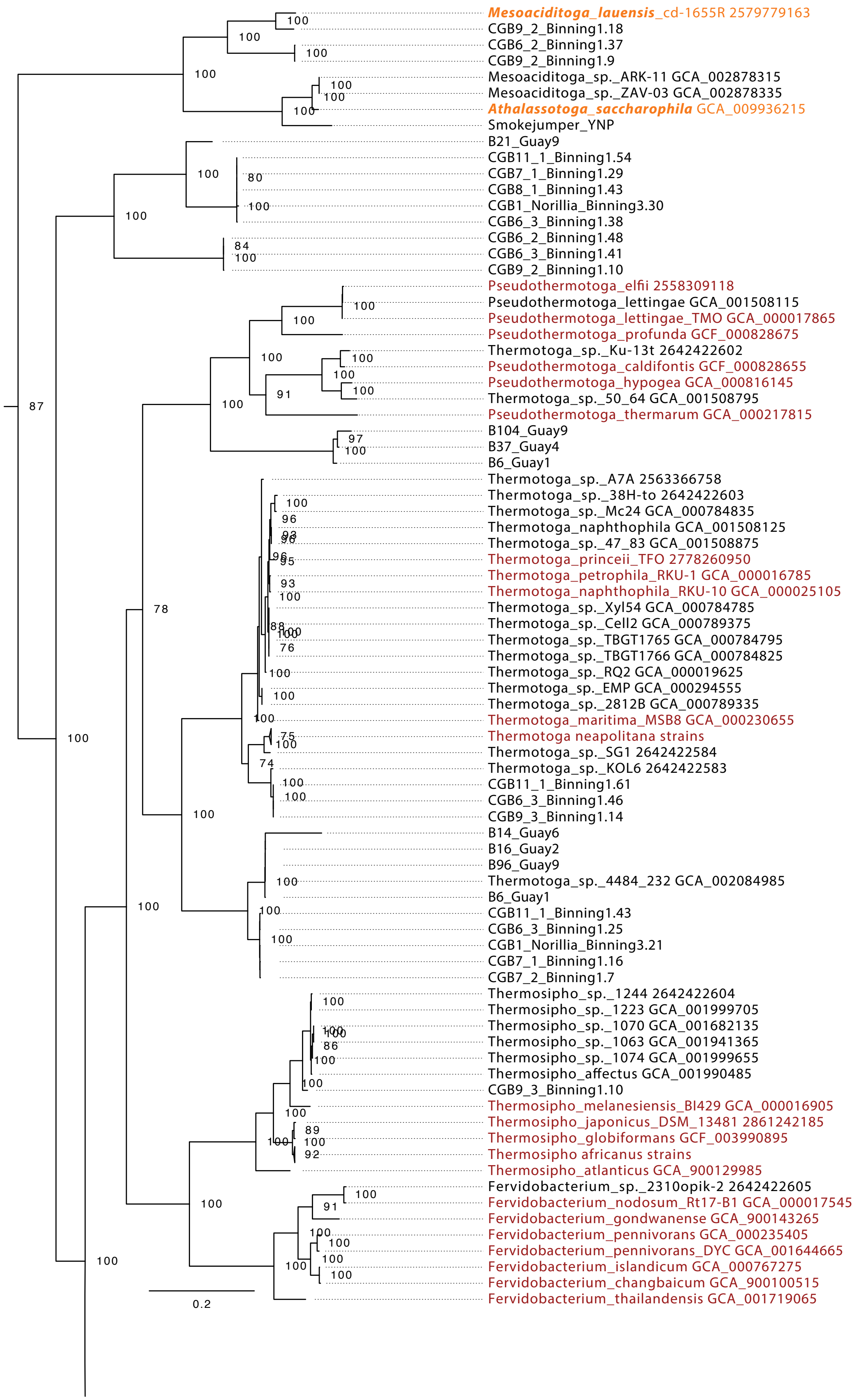
Mesoaciditoga_lauensis_cd-1655R 2579779163
CGB9_2_Binning1.18
CGB6_2_Binning1.37
CGB9_2_Binning1.9
Mesoaciditoga_sp._ARK-11 GCA_002878315
Mesoaciditoga_sp._ZAV-03 GCA_002878335
Athalassotoga_saccharophila GCA_009936215
Smokejumper_YNP
B21_Guay9
CGB11_1_Binning1.54
CGB7_1_Binning1.29
CGB8_1_Binning1.43
CGB1_Norillia_Binning3.30
CGB6_3_Binning1.38
CGB6_2_Binning1.48
CGB6_3_Binning1.41
CGB9_2_Binning1.10
Pseudothermotoga_elfii 2558309118
Pseudothermotoga_lettingae GCA_001508115
Pseudothermotoga_lettingae_TMO GCA_000017865
Pseudothermotoga_profunda GCF_000828675
Thermotoga_sp._Ku-13t 2642422602
Pseudothermotoga_caldifontis GCF_000828655
Pseudothermotoga_hypogea GCA_000816145
Thermotoga_sp._50_64 GCA_001508795
Pseudothermotoga_thermarum GCA_000217815
B104_Guay9
B37_Guay4
B6_Guay1
Thermotoga_sp._A7A 2563366758
Thermotoga_sp._38H-to 2642422603
Thermotoga_sp._Mc24 GCA_000784835
Thermotoga_naphthophila GCA_001508125
Thermotoga_sp._47_83 GCA_001508875
Thermotoga_princeii_TFO 2778260950
Thermotoga_petrophila_RKU-1 GCA_000016785
Thermotoga_naphthophila_RKU-10 GCA_000025105
Thermotoga_sp._Xyl54 GCA_000784785
Thermotoga_sp._Cell2 GCA_000789375
Thermotoga_sp._TBGT1765 GCA_000784795
Thermotoga_sp._TBGT1766 GCA_000784825
Thermotoga_sp._RQ2 GCA_000019625
Thermotoga_sp._EMP GCA_000294555
Thermotoga_sp._2812B GCA_000789335
Thermotoga_maritima_MSB8 GCA_000230655
Thermotoga neapolitana strains
Thermotoga_sp._SG1 2642422584
Thermotoga_sp._KOL6 2642422583
CGB11_1_Binning1.61
CGB6_3_Binning1.46
CGB9_3_Binning1.14
B14_Guay6
B16_Guay2
B96_Guay9
Thermotoga_sp._4484_232 GCA_002084985
B6_Guay1
CGB11_1_Binning1.43
CGB6_3_Binning1.25
CGB1_Norillia_Binning3.21
CGB7_1_Binning1.16
CGB7_2_Binning1.7
Thermosipho_sp._1244 2642422604
Thermosipho_sp._1223 GCA_001999705
Thermosipho_sp._1070 GCA_001682135
Thermosipho_sp._1063 GCA_001941365
Thermosipho_sp._1074 GCA_001999655
Thermosipho_affectus GCA_001990485
CGB9_3_Binning1.10
Thermosipho_melanesiensis_BI429 GCA_000016905
Thermosipho_japonicus_DSM_13481 2861242185
Thermosipho_globiformans GCF_003990895
Thermosipho africanus strains
Thermosipho_atlanticus GCA_900129985
Fervidobacterium_sp._2310opik-2 2642422605
Fervidobacterium_nodosum_Rt17-B1 GCA_000017545
Fervidobacterium_gondwanense GCA_900143265
Fervidobacterium_pennivorans GCA_000235405
Fervidobacterium_pennivorans_DYC GCA_001644665
Fervidobacterium_islandicum GCA_000767275
Fervidobacterium_changbaicum GCA_900100515
Fervidobacterium_thailandensis GCA_001719065
0.2

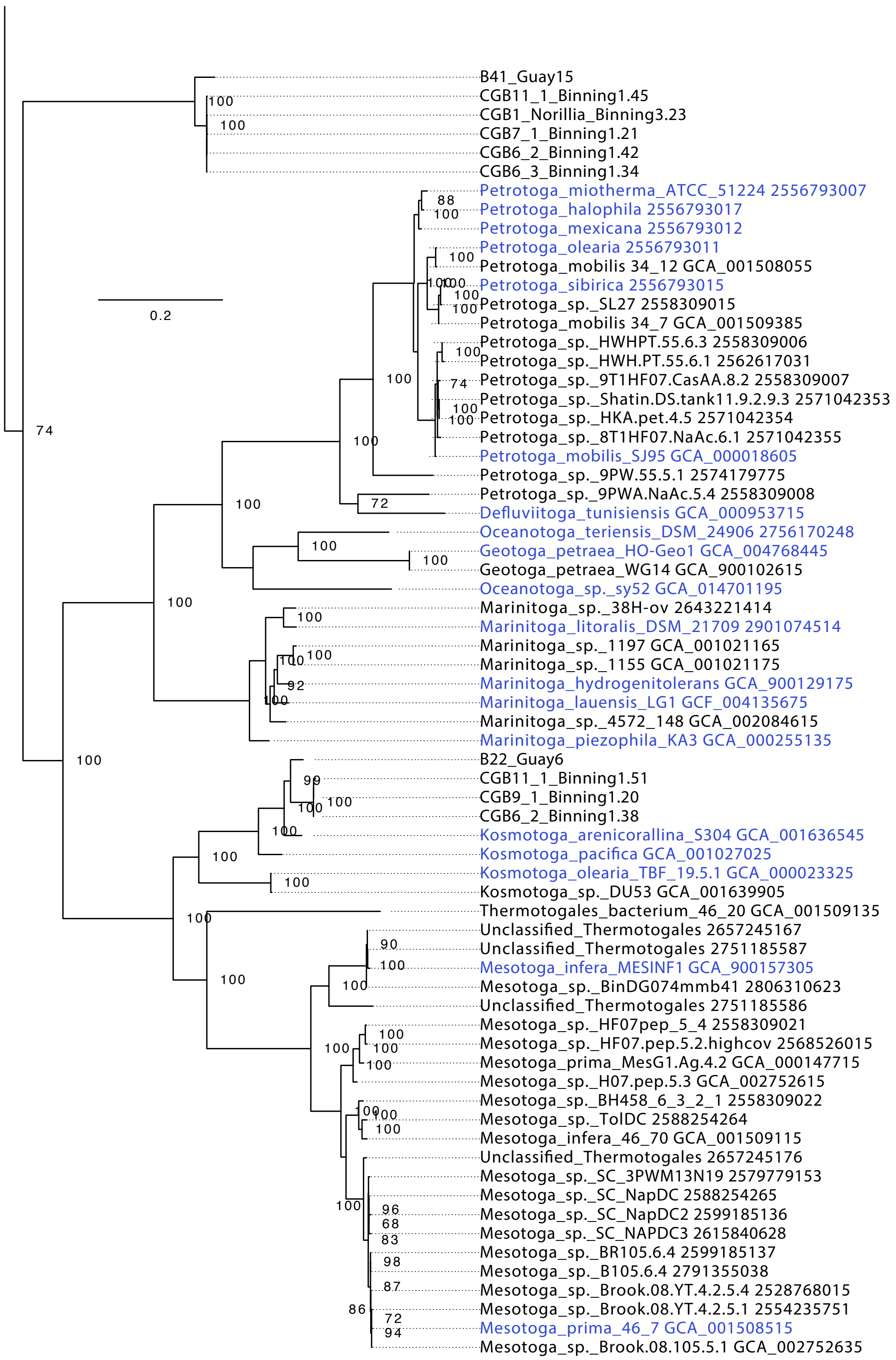
0.2
B41_Guay15
CGB11_1_Binning1.45
CGB1_Norillia_Binning3.23
CGB7_1_Binning1.21
CGB6_2_Binning1.42
CGB6_3_Binning1.34
Petrotoga_miotherma_ATCC_51224 2556793007
Petrotoga_halophila 2556793017
Petrotoga_mexicana 2556793012
Petrotoga_olearia 2556793011
Petrotoga_mobilis 34_12 GCA_001508055
Petrotoga_sibirica 2556793015
Petrotoga_sp._SL27 2558309015
Petrotoga_mobilis 34_7 GCA_001509385
Petrotoga_sp._HWHPT.55.6.3 2558309006
Petrotoga_sp._HWH.PT.55.6.1 2562617031
Petrotoga_sp._9T1HF07.CasAA.8.2 2558309007
Petrotoga_sp._Shatin.DS.tank11.9.2.9.3 2571042353
Petrotoga_sp._HKA.pet.4.5 2571042354
Petrotoga_sp._8T1HF07.NaAc.6.1 2571042355
Petrotoga_mobilis_SJ95 GCA_000018605
Petrotoga_sp._9PW.55.5.1 2574179775
Petrotoga_sp._9PWA.NaAc.5.4 2558309008
Defluviitoga_tunisiensis GCA_000953715
Oceanotoga_teriensis_DSM_24906 2756170248
Geotoga_petraea_HO-Geo1 GCA_004768445
Geotoga_petraea_WG14 GCA_900102615
Oceanotoga_sp._sy52 GCA_014701195
Marinitoga_sp._38H-ov 2643221414
Marinitoga_litoralis_DSM_21709 2901074514
Marinitoga_sp._1197 GCA_001021165
Marinitoga_sp._1155 GCA_001021175
Marinitoga_hydrogenitolerans GCA_900129175
Marinitoga_lauensis_LG1 GCF_004135675
Marinitoga_sp._4572_148 GCA_002084615
Marinitoga_piezophila_KA3 GCA_000255135
B22_Guay6
CGB11_1_Binning1.51
CGB9_1_Binning1.20
CGB6_2_Binning1.38
Kosmotoga_arenicorallina_S304 GCA_001636545
Kosmotoga_pacifica GCA_001027025
Kosmotoga_olearia_TBF_19.5.1 GCA_000023325
Kosmotoga_sp._DU53 GCA_001639905
Thermotogales_bacterium_46_20 GCA_001509135
Unclassified_Thermotogales 2657245167
Unclassified_Thermotogales 2751185587
Mesotoga_infera_MESINF1 GCA_900157305
Mesotoga_sp._BinDG074mmb41 2806310623
Unclassified_Thermotogales 2751185586
Mesotoga_sp._HF07pep_5_4 2558309021
Mesotoga_sp._HF07.pep.5.2.highcov 2568526015
Mesotoga_prima_MesG1.Ag.4.2 GCA_000147715
Mesotoga_sp._H07.pep.5.3 GCA_002752615
Mesotoga_sp._BH458_6_3_2_1 2558309022
Mesotoga_sp._TolDC 2588254264
Mesotoga_infera_46_70 GCA_001509115
Unclassified_Thermotogales 2657245176
Mesotoga_sp._SC_3PWM13N19 2579779153
Mesotoga_sp._SC_NapDC 2588254265
Mesotoga_sp._SC_NapDC2 2599185136
Mesotoga_sp._SC_NAPDC3 2615840628
Mesotoga_sp._BR105.6.4 2599185137
Mesotoga_sp._B105.6.4 2791355038
Mesotoga_sp._Brook.08.YT.4.2.5.4 2528768015
Mesotoga_sp._Brook.08.YT.4.2.5.1 2554235751
Mesotoga_prima_46_7 GCA_001508515
Mesotoga_sp._Brook.08.105.5.1 GCA_002752635

**Supplementary Figure 4. Concatenated ribosomal proteins phylogeny of described *Thermotogota* species and MAGs.** The maximum likelihood tree was reconstructed under the LG+F+R5 substitution model. Bootstrap supports ≥70% are shown. Described species of *Thermotogales*, *Petrotogales*, and *Mesoaciditogales* are shown in red, blue and orange font, respectively. The taxa in black font represent MAGs. Type species of *Mesoaciditogales* are shown in bold font. The tree was rooted using outgroup taxa *Clostridium acetobutylicum* (GCA_000008765), *C. botulinum* (GCA_000063585), *Alkaliphilus metalliredigens* (GCA_000016985), *Bacillus velezensis* (GCA_000875875), *B. cereus* (GCA_003122725), Bacillus_safensis GCA_003097715, *Saccharibacillus sacchari* (GCF_000585395), *Brevibacillus brevis* (GCF_003385915), *Pseudomonas fluorescens* (GCF_003701995), *Thermus thermophilus* (GCA_000091545), *Marinithermus hydrothermalis* (GCA_000195335), *Aquifex aeolicus* (GCF_000008625), *Persephonella marina* (GCF_000021565), *Dictyoglomus thermophilum* (GCF_000020965), *D. turgidum* (GCF_000021645), and MAGs “B3_Guay13”, “CGB8_1_Binning1.39”, “CGB9_2_Binning1.8. The outgroup is not shown. Scale bar, substitutions per site. Tree is available in Newick format in the **FigShare repository** (see **Methods** for details).

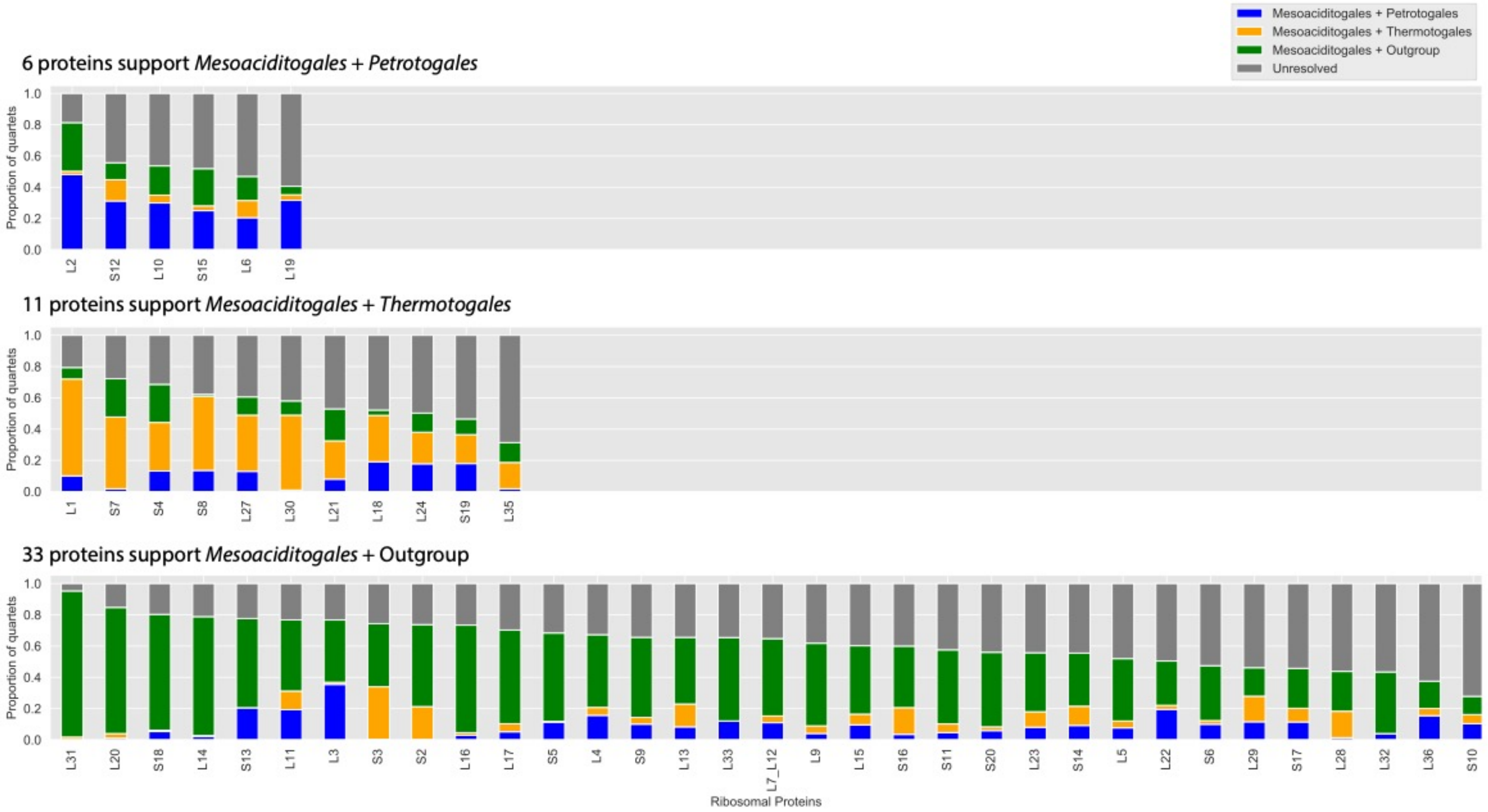


**Supplementary Figure 5. Summary of the likelihood mapping analyses of individual ribosomal proteins**. Each ribosomal protein was analyzed using likelihood mapping, with all possible quartets analyzed. The taxa were divided into four groups: outgroup spp., *Mesoaciditogales* spp., *Petrotogales* spp., and *Thermotogales* spp. The ribosomal proteins are grouped into three panels depending on their support for three possible tree topologies for the four groups. Each bar represents an individual ribosomal protein. The bar is colored by the proportion of quartets (i) strongly supporting each of the three possible topologies (green, yellow, blue), as defined by the basins of attractions $A_1$, $A_2$, and $A_3$ in Strimmer and von Haeseler (1997), and (ii) not strongly supporting any of the three topologies (gray), defined as quartets in simplex areas $A_*$, $A_{13}$, $A_{23}$, and $A_{12}$ (Strimmer and von Haeseler, 1997).

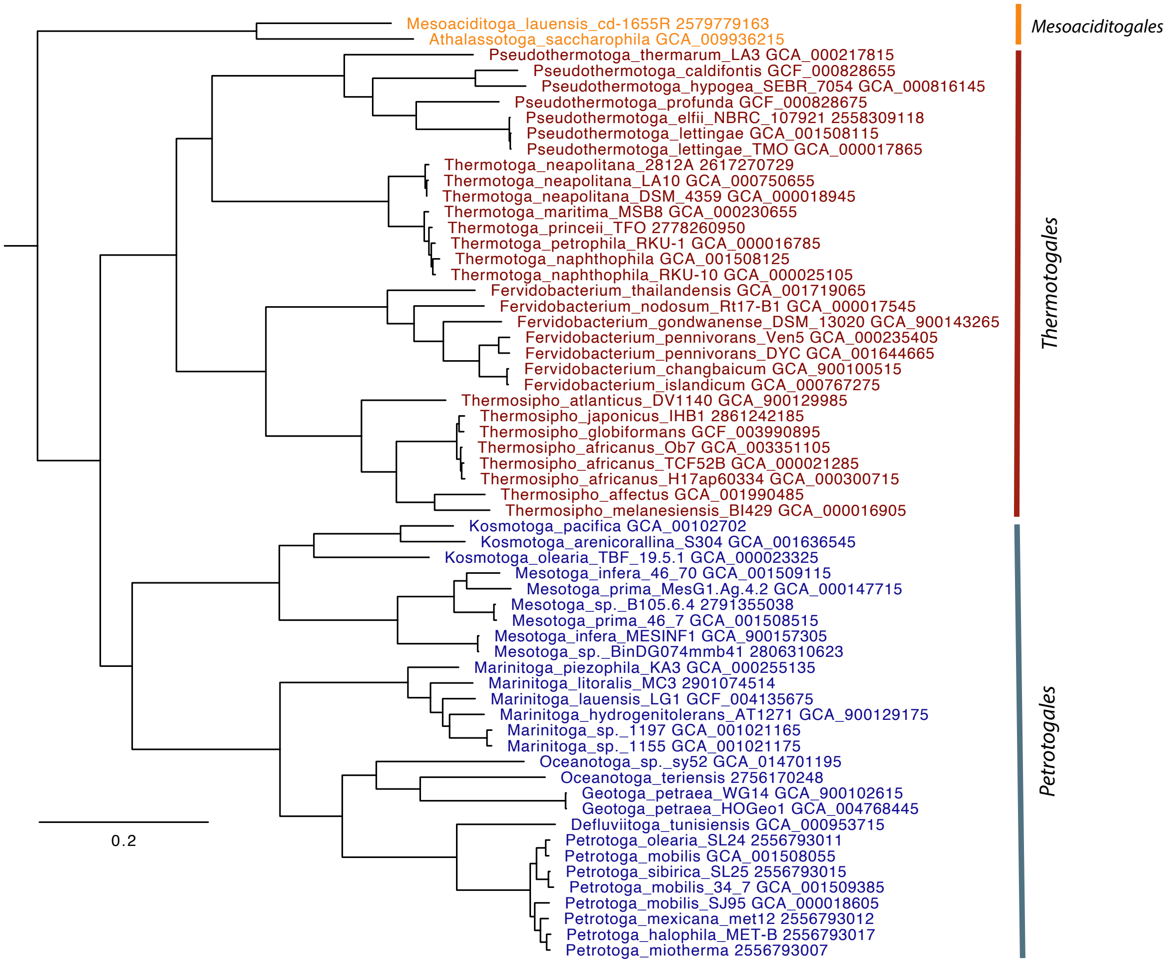


**Supplementary Figure 6. Single-copy core gene tree reconstructed from a recoded dataset**. The SCC genes alignment was recoded into 10 states using the amino acid bins [A],[G],[P],[S],[T],[C], [DENQ], [HKR], [ILMV], and [FWY]. The maximum likelihood phylogeny was built using the GTR+G model for multistate inference. The taxa labels are colored by their order membership: *Petrotogales* are in blue, *Mesoaciditogales* are in orange, and *Thermotogales* are in red. The tree was rooted using outgroup taxa *Clostridium acetobutylicum* (GCA_000008765), *C. botulinum* (GCA_000063585), *Alkaliphilus metalliredigens* (GCA_000016985), *Bacillus velezensis* (GCA_000875875), *B. cereus* (GCA_003122725), Bacillus_safensis GCA_003097715, *Saccharibacillus sacchari* (GCF_000585395), *Brevibacillus brevis* (GCF_003385915), *Pseudomonas fluorescens* (GCF_003701995), *Thermus thermophilus* (GCA_000091545), *Marinithermus hydrothermalis* (GCA_000195335), *Aquifex aeolicus* (GCF_000008625), *Persephonella marina* (GCF_000021565), *Dictyoglomus thermophilum* (GCF_000020965), *D. turgidum* (GCF_000021645). The outgroup is not shown. Scale, substitutions per site. The script used for sequence recoding, the recoded alignment file, and the tree in Newick format are available in the **FigShare repository** (see **Methods** for details).

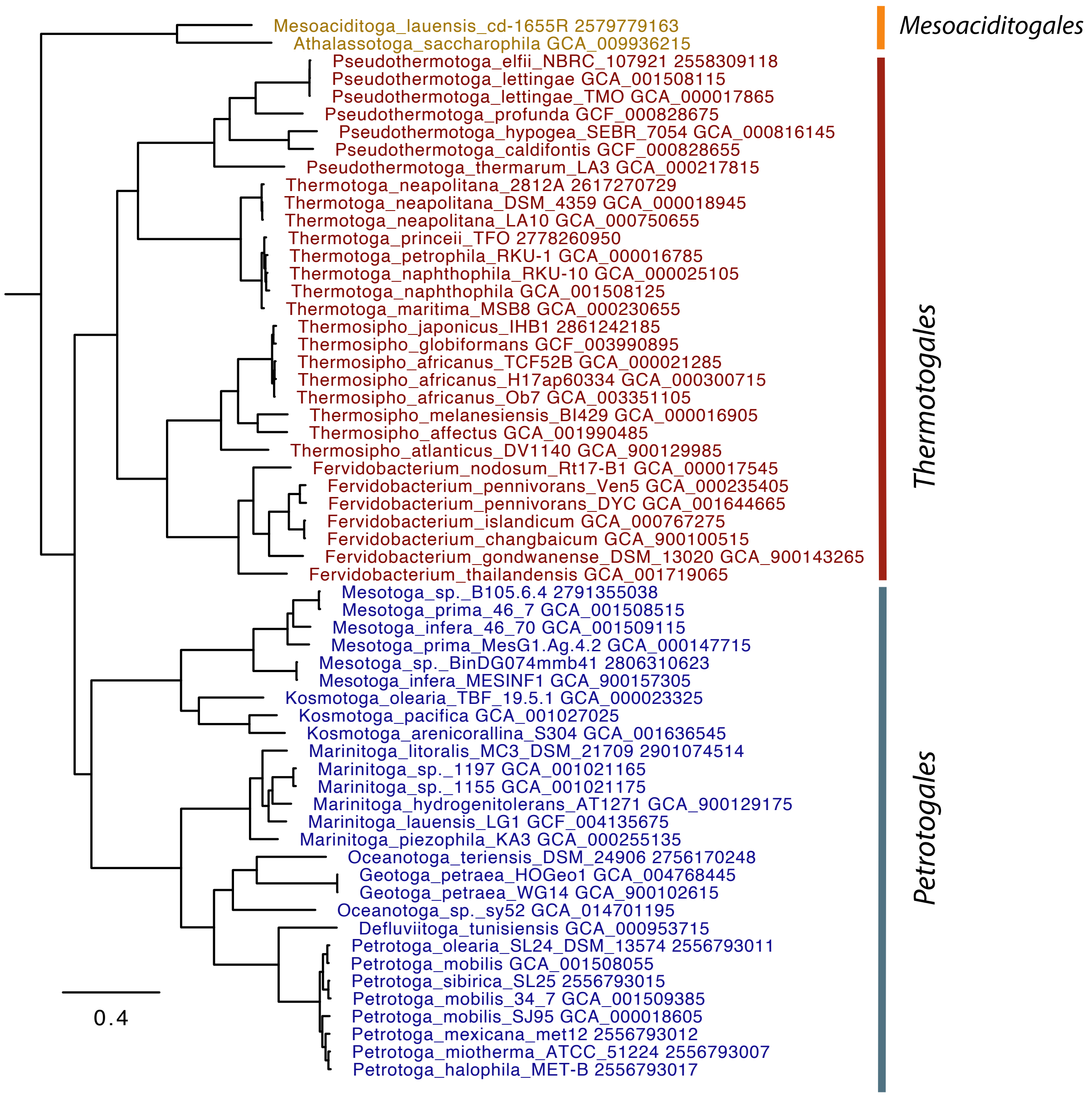


**Supplementary Figure 7. Single-copy core gene tree reconstructed using PMSF model**. The organism names are colored by their order membership: *Petrotogales* are in blue, *Mesoaciditogales* are in orange, and *Thermotogales* are in red. The tree was rooted using outgroup taxa *Clostridium acetobutylicum* (GCA_000008765), *C. botulinum* (GCA_000063585), *Alkaliphilus metalliredigens* (GCA_000016985), *Bacillus velezensis* (GCA_000875875), *B. cereus* (GCA_003122725), Bacillus_safensis GCA_003097715, *Saccharibacillus sacchari* (GCF_000585395), *Brevibacillus brevis* (GCF_003385915), *Pseudomonas fluorescens* (GCF_003701995), *Thermus thermophilus* (GCA_000091545), *Marinithermus hydrothermalis* (GCA_000195335), *Aquifex aeolicus* (GCF_000008625), *Persephonella marina* (GCF_000021565), *Dictyoglomus thermophilum* (GCF_000020965), *D. turgidum* (GCF_000021645). The outgroup is not shown. Scale, substitutions per site. Tree is available in Newick format in the **FigShare repository** (see **Methods** for details).

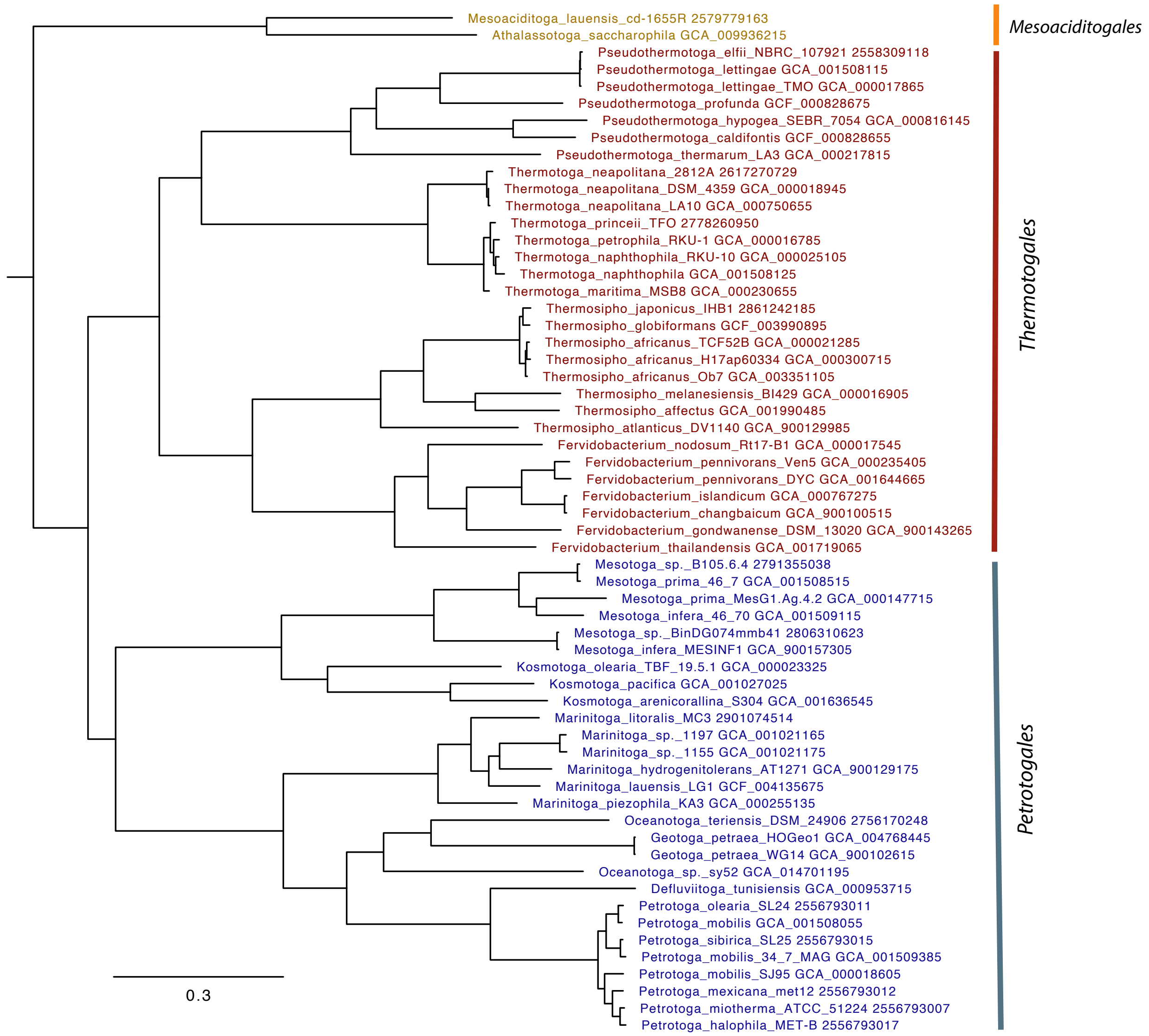


**Supplementary Figure 8. Single-copy core gene tree reconstructed using the GHOST model.** The taxa labels are colored by their order membership: *Petrotogales* are in blue, *Mesoaciditogales* are in orange, and *Thermotogales* are in red. The tree was rooted using outgroup taxa *Clostridium acetobutylicum* (GCA_000008765), *C. botulinum* (GCA_000063585), *Alkaliphilus metalliredigens* (GCA_000016985), *Bacillus velezensis* (GCA_000875875), *B. cereus* (GCA_003122725), Bacillus_safensis GCA_003097715, *Saccharibacillus sacchari* (GCF_000585395), *Brevibacillus brevis* (GCF_003385915), *Pseudomonas fluorescens* (GCF_003701995), *Thermus thermophilus* (GCA_000091545), *Marinithermus hydrothermalis* (GCA_000195335), *Aquifex aeolicus* (GCF_000008625), *Persephonella marina* (GCF_000021565), *Dictyoglomus thermophilum* (GCF_000020965), *D. turgidum* (GCF_000021645). The outgroup is not shown. Scale, substitutions per site. Tree is available in Newick format in the **FigShare repository** (see **Methods** for details).

**Supplementary Table 1. 16S rRNA gene nucleotide sequence identity between individual *Thermotogota* species (first column) and the two *Mesoaciditogales* species (second and third column).** The table cells are color-coded on a gradient from most similar (green) to most different (red). GenBank accession numbers are listed after each taxon name.

| **Described *Thermotogota* species** | ***Athalassotoga saccharophila* AB898788.1** | ***Mesoaciditoga lauensis* cd-1655R NR_125611.1** |
|---|---|---|
| *Athalassotoga saccharophila AB898788.1* | 100.0% | 85.6% |
| *Mesoaciditoga lauensis* cd-1655R NR_125611.1 | 85.6% | 100.0% |
| *Defluviitoga tunisiensis* SulfLac1 NR_122085.1 | 73.0% | 74.3% |
| *Kosmotoga arenicorallina* S304 NR_113012.1 | 76.0% | 78.3% |
| *Kosmotoga olearia* NR_044583.1 | 77.1% | 78.7% |
| *Kosmotoga pacifica* KC119212.1 | 75.7% | 77.8% |
| *Kosmotoga shengliensis* 2SM-2 EU276414.2 | 75.3% | 77.4% |
| *Mesotoga infera* VNs100 NR_118572.1 | 75.9% | 76.5% |
| *Mesotoga prima* NR_102952.1 | 75.9% | 76.8% |
| *Marinitoga camini* NR_028907.1 | 74.8% | 75.3% |
| *Marinitoga hydrogenitolerans* NR_042320.1 | 74.3% | 74.8% |
| *Marinitoga lauensis* LG1 MH457605.1 | 74.5% | 76.1% |
| *Marinitoga litoralis* MC3 NR_104518.1 | 74.4% | 75.1% |
| *Marinitoga okinawensis* NR_041466.1 | 74.4% | 74.7% |
| *Marinitoga piezophila* NR_074102.1 | 75.7% | 75.8% |
| *Marinitoga* sp. 2PyrY55-1 16 KX024844.1 | 75.8% | 76.1% |
| *Geotoga aestuarianus* T3B AF509468.1 | 73.7% | 75.3% |
| *Geotoga petraea* T5 L10658.1 | 72.5% | 74.2% |
| *Geotoga subterranea* CC-1 L10659.1 | 74.2% | 75.0% |
| *Oceanotoga teriensis* OCT74 NR_116378.1 | 71.8% | 73.1% |
| *Tepiditoga spiralis* sy52 LC485113.1 | 74.7% | 77.0% |
| *Petrotoga halophila* MET-B AY800102.1 | 73.6% | 74.1% |
| *Petrotoga japonica* AR80 GQ385331.1 | 72.8% | 73.0% |
| *Petrotoga mexicana* Met-12 AY125964.1 | 73.4% | 73.6% |
| *Petrotoga miotherma* DSM10691 FR733705.1 | 74.2% | 74.3% |
| *Petrotoga mobilis* SJ95 Y15479.1 | 73.9% | 73.8% |
| *Petrotoga olearia* SL24 AJ311703.1 | 73.5% | 73.5% |
| *Petrotoga siberica* SL25 AJ311702.1 | 73.9% | 74.4% |
| *Thermosipho activus* Rift-s3 NR_134221.1 | 74.3% | 76.5% |
| *Thermosipho affectus* NR_117284.1 | 74.3% | 75.2% |
| *Thermosipho africanus* Ob7 DQ647057.1 | 75.2% | 76.6% |
| *Thermosipho atlanticus* NR_029020.1 | 74.9% | 76.0% |
| *Thermosipho ferriphilus* AF491334.1 | 74.1% | 76.8% |
| *Thermosipho geolei* NR_025389.1 | 74.8% | 76.6% |

| | | |
|---|---|---|
| *Thermosipho globiformans* AB257289.1 | 75.8% | 77.2% |
| *Thermosipho japonicus* NR_024726.1 | 74.4% | 76.3% |
| *Thermosipho melanesiensis* NR_102981.1 | 75.9% | 77.3% |
| *Thermotoga maritima* MSB-8 NR_029163.2 | 75.3% | 77.1% |
| *Thermotoga naphthophila* RKU-10 NR_074952.1 | 75.7% | 77.0% |
| *Thermotoga neapolitana* NS-E NR_074959.1 | 75.3% | 76.8% |
| *Thermotoga petrophila* RKU-1 NR_074991.1 | 75.4% | 76.8% |
| *Fervidobacterium changbaicum* NR_043248.1 | 76.8% | 76.7% |
| *Fervidobacterium gondwanense* NR_036997.1 | 76.1% | 76.0% |
| *Fervidobacterium islandicum* NR_044730.2 | 76.0% | 76.0% |
| *Fervidobacterium nodosum* NR_074093.1 | 76.0% | 76.7% |
| *Fervidobacterium pennivorans* NR_074097.1 | 76.4% | 77.9% |
| *Fervidobacterium riparium* NR_108234.1 | 75.7% | 76.1% |
| *Fervidobacterium thailandense* FC2004 KJ473436.1 | 76.3% | 78.8% |
| *Pseudothermotoga caldifontis* AZM44c09 NR_133903.1 | 74.5% | 76.6% |
| *Pseudothermotoga elfii* SERB NR_026201.1 | 74.4% | 75.1% |
| *Pseudothermotoga elfii* subsp. *lettingae* TMO NR_074951.1 | 75.2% | 76.0% |
| *Pseudothermotoga elfii* subsp. *subterranea* SL1T NR_117753.1 | 75.2% | 76.0% |
| *Pseudothermotoga hypogea* NBRC 106472 AP014508.1 | 75.0% | 76.6% |
| *Pseudothermotoga profunda* AZM34c06 NR_133904.1 | 75.2% | 75.9% |
| *Pseudothermotoga thermarum* LA3 NR_074833.1 | 75.4% | 76.0% |
| *Staphylococcus aureus* NR_118997.1 | 71.7% | 72.6% |
| *Thermocrinis ruber* NR_121741.1 | 69.7% | 70.4% |
| *Hydrogenivirga caldilitori* NR_024824.1 | 68.1% | 69.3% |
| *Hydrogenobacter thermophila* NR_074870.1 | 70.5% | 71.4% |
| *Aquifex aeolicus* NR_114796.1 | 68.1% | 69.7% |
| *Coprothermobacter platensis* Y08935.1 | 70.7% | 72.0% |
| *Dictyoglomus thermophilum* NR_029235.1 | 73.4% | 74.7% |
| *Persephonella marina* NR_102828.1 | 71.7% | 71.9% |

**Supplementary Table 2. Summary of 16S rRNA gene's nucleotide identities between members of *Mesoaciditogales* order, two other *Thermotogota* orders and two *Petrotogales* families.** The data is summarized from the individual values listed in Supplementary Table 1.

| ***Mesaoaciditogales* identity to:** | **Mean** | **Median** | **Minimum** | **Maximum** |
|---|---|---|---|---|
| *Petrotogales* order | 74.9% | 74.6% | 71.8% | 78.7% |
| *Thermotogales* order | 75.9% | 76.0% | 74.1% | 78.8% |
| *Kosmotogaceae* family | 76.8% | 76.7% | 75.3% | 78.7% |
| *Petrotogaceae* family | 74.3% | 74.3% | 71.8% | 77.0% |

**Supplementary Table 3. 16S rRNA gene's GC-content of full length sequences and of predicted stem regions only in *Thermotogota* species**. The organism names are colored by their order membership: *Petrotogales* are in blue, *Mesoaciditogales* are in orange, and *Thermotogales* are in red.

| NCBI Nucleotide Accession | Organism Name | 16S rRNA full length GC, % | 16S rRNA stem region GC, % |
|---|---|---|---|
| AF509468.1 | *Geotoga aestuarianus T3B* | 39.9 | 65.7 |
| NR_104518.1 | *Marinitoga litoralis MC3* | 42.2 | 68.1 |
| NR_116378.1 | *Oceanotoga teriensis OCT74* | 42.8 | 64.9 |
| AY800102.1 | *Petrotoga halophila MET-B* | 43.5 | 64.9 |
| Y15479.1 | *Petrotoga mobilis SJ95* | 43.7 | 65.1 |
| GQ385331.1 | *Petrotoga japonica AR80* | 43.9 | 65.5 |
| AJ311702.1 | *Petrotoga siberica SL25* | 44.0 | 65.5 |
| AJ311703.1 | *Petrotoga olearia SL24* | 44.0 | 65.5 |
| LC485113.1 | *Tepiditoga spiralis sy52* | 44.1 | 64.1 |
| NR_122085.1 | *Defluviitoga tunisiensis SulfLac1* | 44.4 | 66.7 |
| FR733705.1 | *Petrotoga miotherma 42-6* | 44.5 | 65.4 |
| AY125964.1 | *Petrotoga mexicana Met-12* | 44.7 | 65.5 |
| MH457605.1 | *Marinitoga lauensis LG1* | 45.2 | 69.1 |
| KJ473436.1 | *Fervidobacterium thailandense FC2004* | 45.3 | 76.1 |
| NR_028907.1 | *Marinitoga camini MV1075* | 45.4 | 68.2 |
| L10658.1 | *Geotoga petraea T5* | 45.8 | 67.4 |
| AB898788.1 | *Athalassotoga saccharophila NAS-01* | 45.8 | 66.8 |
| NR_125611.1 | *Mesoaciditoga lauensis cd-1655R* | 45.9 | 70.3 |
| NR_118572.1 | *Mesotoga infera VNs100* | 45.9 | 67.2 |
| DQ647057.1 | *Thermosipho africanus Ob7* | 46.0 | 74.5 |
| NR_041466.1 | *Marinitoga okinawensis TFS10-5* | 46.1 | 69.0 |
| NR_042320.1 | *Marinitoga hydrogenitolerans AT1271* | 46.4 | 68.4 |
| NR_102952.1 | *Mesotoga prima MesG1.Ag.4.2* | 46.6 | 68.4 |
| AB257289.1 | *Thermosipho globiformans MN14* | 47.0 | 74.0 |
| KX024844.1 | *Marinitoga arctica 2PyrY55-1* | 47.1 | 70.0 |
| L10659.1 | *Geotoga subterranea CC-1* | 47.1 | 67.2 |
| NR_113012.1 | *Kosmotoga arenicorallina S304* | 47.7 | 73.2 |
| NR_044730.2 | *Fervidobacterium islandicum H-21* | 48.3 | 73.7 |
| NR_036997.1 | *Fervidobacterium gondwanense AB39* | 48.3 | 72.0 |
| MK818504.1 | *Thermosipho ferrireducens JL129W03* | 48.3 | 75.0 |
| NR_108234.1 | *Fervidobacterium riparium 1445t* | 48.7 | 72.4 |
| NR_074102.1 | *Marinitoga piezophila KA3* | 48.8 | 70.3 |
| EU276414.2 | *Kosmotoga shengliensis 2SM-2* | 48.9 | 75.0 |

| | | | |
|---|---|---|---|
| NR_074093.1 | *Fervidobacterium nodosum Rt17-B1* | 49.3 | 73.3 |
| NR_026201.1 | *Pseudothermotoga elfii SERB* | 49.4 | 74.0 |
| KC119212.1 | *Kosmotoga pacifica SLHLJ1* | 49.5 | 73.7 |
| NR_029020.1 | *Thermosipho atlanticus DV1140* | 49.9 | 75.0 |
| NR_117284.1 | *Thermosipho affectus ik275mar* | 50.0 | 74.5 |
| NR_024726.1 | *Thermosipho japonicus IHB1* | 50.0 | 74.3 |
| NR_025389.1 | *Thermosipho geolei SL31 DSM 13256* | 50.1 | 74.1 |
| NR_043248.1 | *Fervidobacterium changbaicum CBS-1* | 50.2 | 73.9 |
| NR_074097.1 | *Fervidobacterium pennivorans Ven5* | 50.2 | 74.8 |
| NR_133904.1 | *Pseudothermotoga profunda AZM34c06* | 50.3 | 74.0 |
| NR_134221.1 | *Thermosipho activus Rift-s3* | 50.5 | 74.6 |
| AF491334.1 | *Thermosipho ferriphilus GB21* | 50.6 | 74.2 |
| NR_117753.1 | *Pseudothermotoga elfii subterranea SL1* | 50.8 | 74.8 |
| NR_074951.1 | *Pseudothermotoga elfii lettingae TMO* | 51.0 | 74.9 |
| NR_102981.1 | *Thermosipho melanesiensis BI429* | 51.4 | 74.7 |
| AP014508.1 | *Pseudothermotoga hypogea SEBR 7054* | 51.7 | 77.0 |
| NR_044583.1 | *Kosmotoga olearia TBF 19.5.1* | 51.9 | 75.1 |
| NR_074833.1 | *Pseudothermotoga thermarum LA3* | 52.7 | 77.5 |
| NR_133903.1 | *Pseudothermotoga caldifontis AZM44c09* | 52.9 | 77.1 |
| NR_074952.1 | *Thermotoga naphthophila RKU-10* | 53.8 | 78.7 |
| NR_074959.1 | *Thermotoga neapolitana NS-E* | 54.2 | 79.1 |
| NR_074991.1 | *Thermotoga petrophila RKU-1* | 54.3 | 78.9 |
| NR_029163.2 | *Thermotoga maritima MSB-8* | 54.3 | 79.1 |

**Supplementary Table 4. Summary of 16S rRNA gene's GC-content within three *Thermotogota* orders and two *Petrotogales*' families.** The data is summarized from the individual values listed in Supplementary Table 3.

| | Full Length | | | |
|---|---|---|---|---|
| **Order/Family** | **Mean, %** | **Median, %** | **Minimum, %** | **Maximum, %** |
| *Thermotogales* order | 50.3 | 50.2 | 45.3 | 54.3 |
| *Petrotogales* order | 45.5 | 45.3 | 39.9 | 51.9 |
| *Kosmotogacae* family | 48.4 | 48.3 | 51.9 | 45.9 |
| *Petrotogaceae* family | 44.7 | 44.4 | 48.8 | 39.9 |
| *Mesociditogales* order | 45.9 | 45.9 | 45.8 | 45.9 |
| | **Stem Regions** | | | |
| **Order/Family** | **Mean, %** | **Median, %** | **Minimum, %** | **Maximum, %** |
| *Thermotogales* order | 75.2 | 74.6 | 72.0 | 79.1 |
| *Petrotogales* order | 68.0 | 67.3 | 64.1 | 75.1 |
| *Kosmotogacae* family | 72.1 | 73.4 | 67.2 | 75.1 |
| *Petrotogaceae* family | 66.6 | 65.7 | 64.1 | 70.0 |
| *Mesoaciditogales* order | 68.6 | 68.3 | 66.8 | 70.3 |
